# Comparative molecular mapping reveals molecular gradients and spatially organized populations in the subthalamic and para-subthalamic regions

**DOI:** 10.1101/2025.05.21.655268

**Authors:** Sylvie Dumas, Chantal Francois, Marie Biviano Rosenkilde, Julia Staryk, Urmas Roostalu, Andreas Heuer, Carine Karachi, Åsa Wallén-Mackenzie

## Abstract

The subthalamic nucleus (STN) is a major target for deep brain stimulation (DBS) in Parkinson’s disease (PD), yet its molecular organization and relationship to the adjacent para-subthalamic nucleus (pSTN) remain incompletely defined. Here, we performed comparative molecular mapping of the STN and pSTN in mouse and macaque, complemented by a human tissue sample, three-dimensional imaging, and publicly available spatial transcriptomic datasets. The STN showed a largely conserved molecular framework across species with gradients across rostrocaudal and dorsolateral axes. However, *HTR2C* was found enriched in the ventral STN of macaque and human tissue samples, while broadly distributed across the mouse STN. *HTR2C* showed opposite gradient of *PVALB*. In the mouse STN, *Nts* expression revealed an additional mediolateral gradient. Publicly available transcriptomic datasets showed concordant expression patterns. In both species, *TAC1* characterized the canonical pSTN but also an extended population reaching dorsally along the medial aspect of the STN. *TAC1*-positive cells were thus found in the region traversed by tyrosine hydroxylase-positive fibers of the medial forebrain bundle (MFB). Together, these findings reveal a spatially heterogeneous, multiaxial molecular organization of the STN and a spatially complex molecular organization of its neighboring brain areas, refining the molecular and anatomical framework of the subthalamic region.

## Introduction

The subthalamic nucleus (STN) is an excitatory component of the basal ganglia and an established target for deep brain stimulation (DBS) in Parkinson’s disease (PD), particularly for the treatment of motor symptoms^1–3^. In PD, abnormal STN activity and altered basal ganglia network dynamics are associated with motor dysfunction, and STN-DBS is thought to alleviate symptoms by modulating pathological activity within these circuits^4,5^. Although stimulation is generally directed toward the dorsolateral, sensorimotor STN, the volume of tissue activated may extend into neighboring or functionally distinct territories^6,7^. STN-DBS can also influence non-motor symptoms, with effects varying across symptom domains and stimulation location^8^ and the STN is targeted in other neurological and neuropsychiatric disorders, including dystonia, Tourette syndrome, and obsessive-compulsive disorder^9–11^. Molecularly defined populations provide an independent framework for investigating this organization and for distinguishing the STN from neighboring structures that may be engaged by stimulation.

Anatomical tracing and neuroimaging studies in primates have described broad sensorimotor, associative, and limbic territories distributed along the dorsoventral to ventromedial axes^12–14^. Such organization into three anatomical-functional territories within the primate STN is commonly represented by the so called tripartite model^12,15,16^ although whether these territories constitute discrete subdivisions or overlapping functional gradients remains unresolved^17–19^. Molecularly defined neuronal populations could therefore provide an independent framework for determining whether proposed anatomical-functional territories correspond to discrete cellular domains or broader molecular gradients.

The molecular landscape of the STN has increasingly been characterized, with recent reviews highlighting its neurochemical composition and emerging molecular organization^20,21^. Among the molecular features contributing to this landscape are calcium-binding proteins such as the parvalbumin protein (PV, encoded by *PVALB*), whose presence in the dorsal STN territory of the primate STN has been recognized^20^.

The STN is predominantly glutamatergic, with GABAergic neurons representing a minor population^22,23^. The vesicular glutamate transporter *VGLUT2* (*SLC17A6/Vglut2*) is expressed by STN neurons and serves as a marker of its glutamatergic population^24–26^, while the homeobox transcription factor *Pitx2* mRNA provides a complementary molecular marker mainly studied in mice where it plays a role in STN development^27,28^. Our previous work demonstrated substantial overlap of *Pitx2* and *Vglut2* expression in the mouse STN and showed that manipulation of this population using mouse genetics, including conditional knockout and optogenetics methodology, affects locomotor behavior and affective function^29–32^. In addition,these findings established *Pitx2* as a genetic access point for the *Vglut2*-positive STN population.

We subsequently used *Pitx2*-targeted single-nucleus RNA sequencing (snRNA-seq) and fluorescent *in situ* hybridization (FISH) to characterize molecular heterogeneity within the mouse STN^33^. *Htr2c*, *Nxph1*, *Nxph4*, and *Kcnab3* showed broad STN expression, whereas *Col24a1* and *Pvalb* displayed opposing dorsoventral distributions. FISH mapping of these markers subdivided the *Vglut2*/*Pitx2*-positive STN into three spatio-molecular domains: a ventral *Col24a1*^+^/*Pvalb*^−^ domain (STN_a_), a central *Col24a1*^+^/*Pvalb^+^*domain (STN_b_), and a dorsal *Col24a1^−^/Pvalb^+^*domain (STN_c_). The dorsal *Pvalb*-positive domain was consistent with the reported distribution of the corresponding PV protein in the primate STN. In a recent comparative analysis, we demonstrated substantial overlap between *PITX2* and *VGLUT2* throughout the macaque STN structure and also verified the presence of *PVALB* mRNA in the dorsal STN of the macaque^32^. These findings prompt the question of whether also additional spatio-molecular features recently identified in the mouse STN ^33^ are conserved across species.

Molecular approaches have also begun to distinguish the STN from the adjacent para-subthalamic nucleus (pSTN), a crescent-shaped region implicated in autonomic, interoceptive, and motivated behaviors^34^. Its boundary with the STN is poorly defined by conventional cytoarchitecture, making molecular markers particularly useful for delineating these neighboring structures. The rat pSTN was described as a molecularly distinct cell group enriched in β-preprotachykinin (*Tac1*) mRNA^35^, and *Tac1* has subsequently been used as a pSTN marker in mice and rats by several laboratories^32,34,36–40^.

Our transcriptomic and FISH analyses identified *Tac1*, *Baiap3*, and *Glra3* as enriched in the pSTN but absent from the STN, and also identified the same enrichment of *TAC1* in the macaque STN, thus establishing *TAC1* as a marker also for the macaque pSTN^32,33^. Other genes, including *Crh*, *Nts*, *Penk*, and *Calb2*, have also been reported as enriched in the mouse pSTN^33,34^, supporting molecular heterogeneity within this region. However, the broader organization and spatial extent of the pSTN across species, and its relationship to the adjacent STN, remain incompletely defined.

Together, these studies point to an emerging molecular framework in which the STN and pSTN comprise spatially organized and heterogeneous neuronal populations, extending beyond the traditional description of the STN as a relatively uniform glutamatergic nucleus and the pSTN as a poorly demarcated neighboring region.

Here, we performed comparative molecular mapping of the STN and pSTN in mouse and macaque using FISH, complemented by one human tissue sample. Recently published public spatial transcriptomic datasets were queried for validation where applicable. Because the spatial relationship between molecularly defined populations and adjacent fiber systems cannot always be resolved from two-dimensional sections alone, we additionally incorporated three-dimensional brain imaging and rendering to examine the anatomical relationship of the STN and pSTN to the medial forebrain bundle (MFB) which also occupies the region. We found that the molecular framework of the STN was broadly conserved between mouse and macaque, while identifying a prominent *HTR2C*-enriched ventral domain in the macaque STN that was also observed in the limited human tissue. In addition to the canonical pSTN, we identified an unexpected *TAC1*-positive population reaching dorsally along the medial aspect of the STN, where it was closely associated with TH-positive fibers of the MFB. Together, these findings extend the emerging molecular framework of subthalamic organization across species and establish molecular landmarks within the STN and neighboring pSTN that provide a spatial basis for understanding their anatomical and functional organization and its relevance to therapeutic neuromodulation.

## Results

### Molecular and anatomical organization of the STN - pSTN region across species

The anatomical-functional model of the primate STN (**Fig. 1a**) and the spatio-molecular model of the mouse STN (**Fig. 1b**) provided the basis for a molecular - anatomical comparative analysis of the STN and pSTN in mouse and macaque across all axes (**Fig. 1c**)

**Figure 1.**
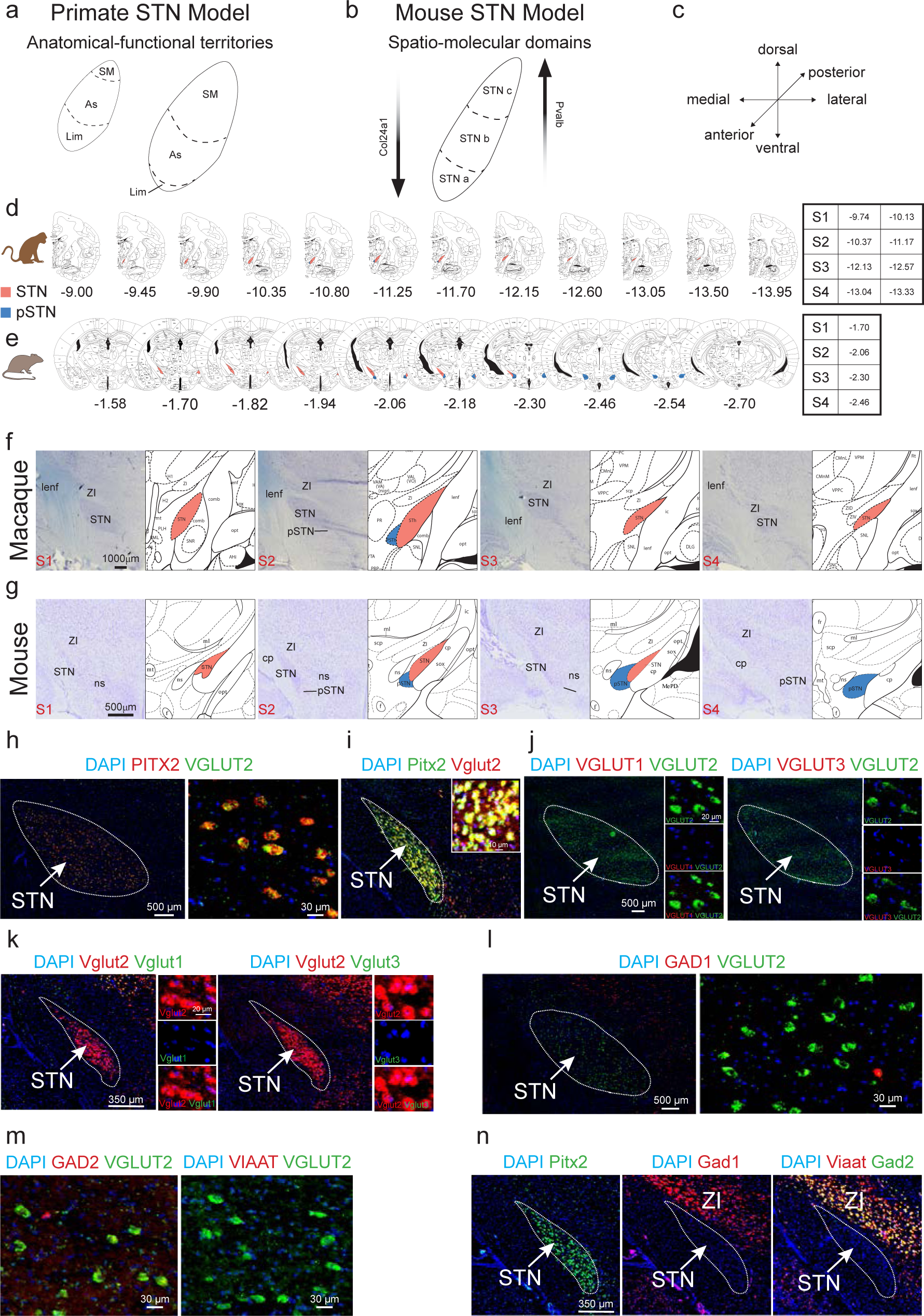
Molecular and anatomical organization of the STN - pSTN region across species. **(a,b)** Schematic representations of the primate tripartite STN model (a) and the mouse spatiomolecular STN model (b). **(c)** Schematic representation of the anatomical section planes across the three axes. **(d,e)** Macaque (d) and mouse (e) brain atlas sections showing the approximate anteroposterior extent of the STN and canonical pSTN. In macaque, the STN extends approximately from Bregma −9.45 to −13.50 mm, with the canonical pSTN located approximately between −10.80 and −11.70 mm. In mouse, the STN extends approximately from Bregma −1.70 to −2.30 mm, with the canonical pSTN located approximately between −2.06 and −2.54 mm. Approximate section levels (S1 - S4) used throughout the study and their corresponding Bregma coordinates are indicated in the tables (anterior to posterior range shown for macaque at each indicated Bregma) **(f,g)** Cresyl violet-stained sections across S1 - S4 matched to corresponding atlas levels in macaque (f) and mouse (g) with approximate Bregma shown in table (for macaque, this is presented as anterior – posterior range; see Methods for complete list); STN is shown in pink and pSTN in blue. **(h - n)** FISH analysis of molecular markers in macaque and mouse STN/pSTN. Probe colors indicate fluorophores; single labeling is shown in red or green and co-localization in yellow. DAPI nuclear counterstaining is shown in blue. Enlarged views show selected regions. **(h)** Macaque STN at S3, *PITX2/VGLUT2*. **(i)** Mouse STN at S2, *Pitx2/Vglut2*. **(j)** Macaque STN, *VGLUT2/VGLUT2* and *VGLUT3/VGLUT2*. **(k)** Mouse STN, *Vglut2/Vglut1* and *Vglut2/Vglut3*. **(l)** Macaque STN, *GAD1/VGLUT2*. **(m)** Macaque STN, *GAD2/VGLUT2* and *VIAAT/VGLUT2*. **(n)** Mouse STN, *Pitx2*, *Gad1* and *Viaat/Gad2*. Scale bars, µm; individual values are indicated in each panel. Full mRNA names and probe information are provided in **Supplementary Table 2**. Abbreviations: cp, cerebral peduncle; FISH, fluorescent *in situ hybridization*; lenf, lenticular fasciculus; ns, nigrostriatal pathway; pSTN, para-subthalamic nucleus; STN, subthalamic nucleus; ZI, zona incerta. Atlas abbreviations follow the original publications, macaque^41^, mouse^42^. S1 - S4 refers to section (S) levels corresponding to approximate Bregma level as specified in d and e.

We first compared the anatomical extent and molecular organization of the STN - pSTN region in mouse and macaque using published anatomical atlases, Nissl staining, and FISH for glutamatergic and inhibitory markers (**Fig. 1**; **Supplementary Figure 1**).

Atlas-defined boundaries differed between species. In macaque, the STN extended approximately from Bregma −9.45 to −13.50 mm, with the pSTN located between approximately −10.80 and −11.70 mm (**Fig. 1d**; **Supplementary Figure 1**)^41^. In mouse, the STN extended approximately from Bregma −1.70 to −2.30 mm, whereas the pSTN occupied a more posterior territory, approximately −2.06 to −2.54 mm (**Fig. 1e**; **Supplementary Figure 1**)^42^.

Three-dimensional rendering of atlas coordinates further illustrated the spatial relationship between the STN and pSTN in the two species (**Supplementary Figure 1**). 3D reconstructions visualized the pSTN as a smaller, elongated structure directly adjoining the ventral aspect of the STN, in agreement with published atlases. The two nuclei displayed close spatial alignment.

Nissl staining identified the expected cellular territory of the STN in both species, whereas the lower cellular density of the pSTN made its boundaries less distinct (**Fig. 1f,g**). FISH using probes to detect glutamatergic and GABAergic neurotransmitter identity was next implemented. *VGLUT2 and PITX2* FISH were used as reference for both macaque and mouse STN and showed substantial overlap in accordance with our previous report ^32^ (**Fig. 1h, i**). Expression patterns of *VGLUT2* and *PITX2* in macaque and human STN were also observed in images available in the Allen Brain Cell (ABC) Atlas (**Supplementary Figure 2, 3**) (https://knowledge.brain-map.org/abcatlas) with a seemingly similar distribution as observed in our material.

*VGLUT1*, *VGLUT3*, *GAD1*, *GAD2* and *VIAAT* were absent or detected only rarely in STN cells in either species (**Fig. 1j-n**).

The section plane of the macaque sections was further validated by comparing *VGLUT2*-positive (e.g. STN and pSTN) areas with neighboring *TH*- and *GAD1*-positive structures. Across the sampled rostrocaudal levels, the relative positions of the neighboring structures corresponded to the atlas-defined anatomy at the coronal section plane (**Supplementary Figure 4**).

### Molecular delineation of *TAC1*-positive canonical pSTN identifies an extended *TAC1*-labeled population in the macaque brain

Having anatomically validated our section material, we next examined molecular markers previously identified as differentially expressed between the STN and pSTN in the mouse. *VGLUT2* FISH was used as reference. *TAC1* FISH identified the canonical pSTN territory and distinguished it from the STN (**Fig. 2a**). *TAC1*-positive cells were concentrated at anterior STN levels, approximately between Bregma −9.94 and −11.77 mm, and were not detected at the more posterior levels examined (**Fig. 2a**) in accordance with atlas, as represented in *Fig. 1* above.

**Figure 2.**
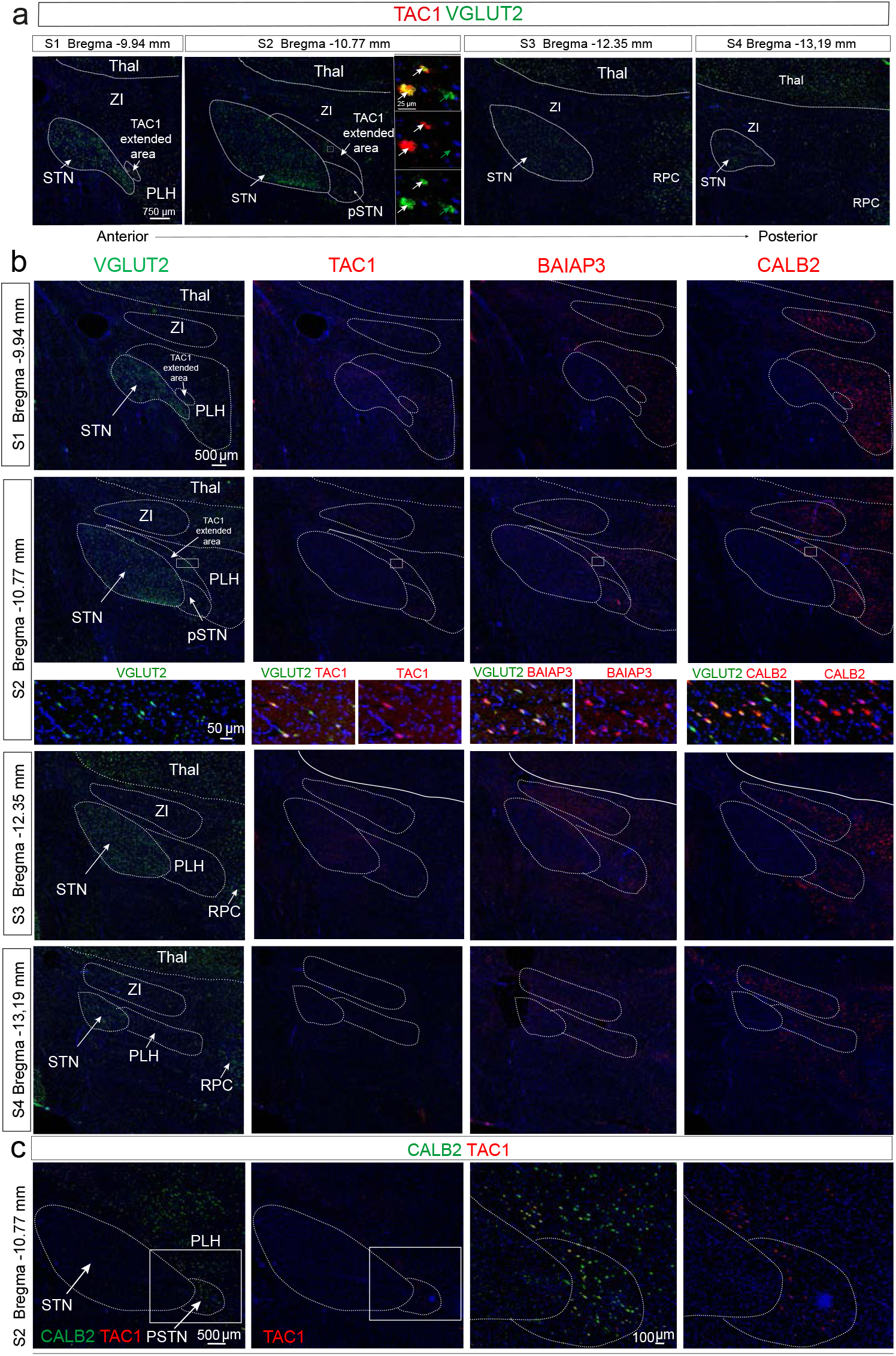
Molecular delineation of the canonical pSTN identifies an extended TAC1-positive area in macaque. FISH analysis of *TAC1* and associated molecular markers in the macaque STN - pSTN region across section levels S1 - S4. The canonical pSTN and a dorsomedial TAC1-positive area adjacent to the canonical pSTN, referred to as the TAC1 extended area, are detected at S1 - S2 but are not detected at S3 - S4. Probe colors indicate fluorophores; single labeling is shown in red or green and co-localization in yellow. Enlarged views show selected regions. **(a)** *TAC1/VGLUT2* across S1 - S4. Arrows indicate the *TAC1* extended area at the medial aspect of the STN, dorsal to the canonical pSTN. **(b)** *VGLUT2*, *TAC1*, *BAIAP3* and *CALB2* at S2. *BAIAP3* and *CALB2* are detected in the canonical pSTN and in *TAC1* extended area and extend laterally into the posterior lateral hypothalamus (PLH). Enlarged views correspond to the regions indicated by squares. **(c)** *CALB2/TAC1* at S2, showing the canonical pSTN and *TAC1* extended area; *CALB2* extends further laterally into the PLH. Scale bars, µm; individual values are indicated in each panel. Bregma coordinates and section levels are indicated. Full mRNA names and probe information are provided in **Supplementary Table 2**. Abbreviations: FISH, fluorescent *in situ* hybridization; pSTN, para-subthalamic nucleus; PLH, posterior lateral hypothalamus; STN, subthalamic nucleus; Thal, thalamus; ZI, zona incerta. S1 - S4 refers to section (S) levels corresponding to approximate Bregma level as specified in Fig. 1d and e.

In addition to the canonical pSTN territory, *TAC1*-positive cells extended dorsally along the medial aspect of the molecularly defined STN and approached its dorsal extent at approximately Bregma −10.77 mm (S2) (**Fig. 2a, b**). *BAIAP3* and *CALB2* showed similar distributions while not completely overlapping with *TAC1* both within the canonical pSTN and within the spatially extended *TAC1*-positive territory; however with these two additional markers extended further laterally into posterior lateral hypothalamus than *TAC1* (**Fig. 2b**). *CALB2* labeling was partially co-localized with *TAC1* in both canonical pSTN and the dorsally extending *TAC1* area (**Fig. 2b, c**)

Because the available anatomical atlases do not establish whether this additional dorsally extending *TAC1*-labeled population represents an extension of the canonical pSTN, we refer to it hereafter as the “extended *TAC1* area” for simplification and for distinguishing it from the canonical atlas-defined pSTN.

Publicly available datasets were queried for validation. Notably, the pSTN is not an annotated structure in the macaque and human brain in the ABC Atlas (https://knowledge.brain-map.org/abcatlas). However, *TAC1* was a searchable mRNA. Images showed *TAC1* positive signal in the area putatively corresponding to the canonical pSTN (the anatomical location ventral to the ventral STN) in macaque, while *TAC1* was not observed in human sample. Further, section level information was not provided in the human dataset. (**Supplementary Figures 2, 3**).

### Several markers originally identified in mouse STN conserved in macaque STN

Having identified the border between STN and pSTN in macaque, we next examined the STN itself with markers previously identified in the mouse STN. *VGLUT2* was used as reference. *NXPH4*, *NXPH1* and *KCNAB3* markers were broadly distributed throughout the *VGLUT2*-positive STN and were largely absent from the adjacent pSTN and the extended *TAC1* area (**Fig. 3a**). Within the STN, their distributions closely resembled that of *VGLUT2*, and showed extensive co-localization with *VGLUT2* within the STN across the anteroposterior axis (S1 - S4) (**Fig. 3b**).

**Figure 3.**
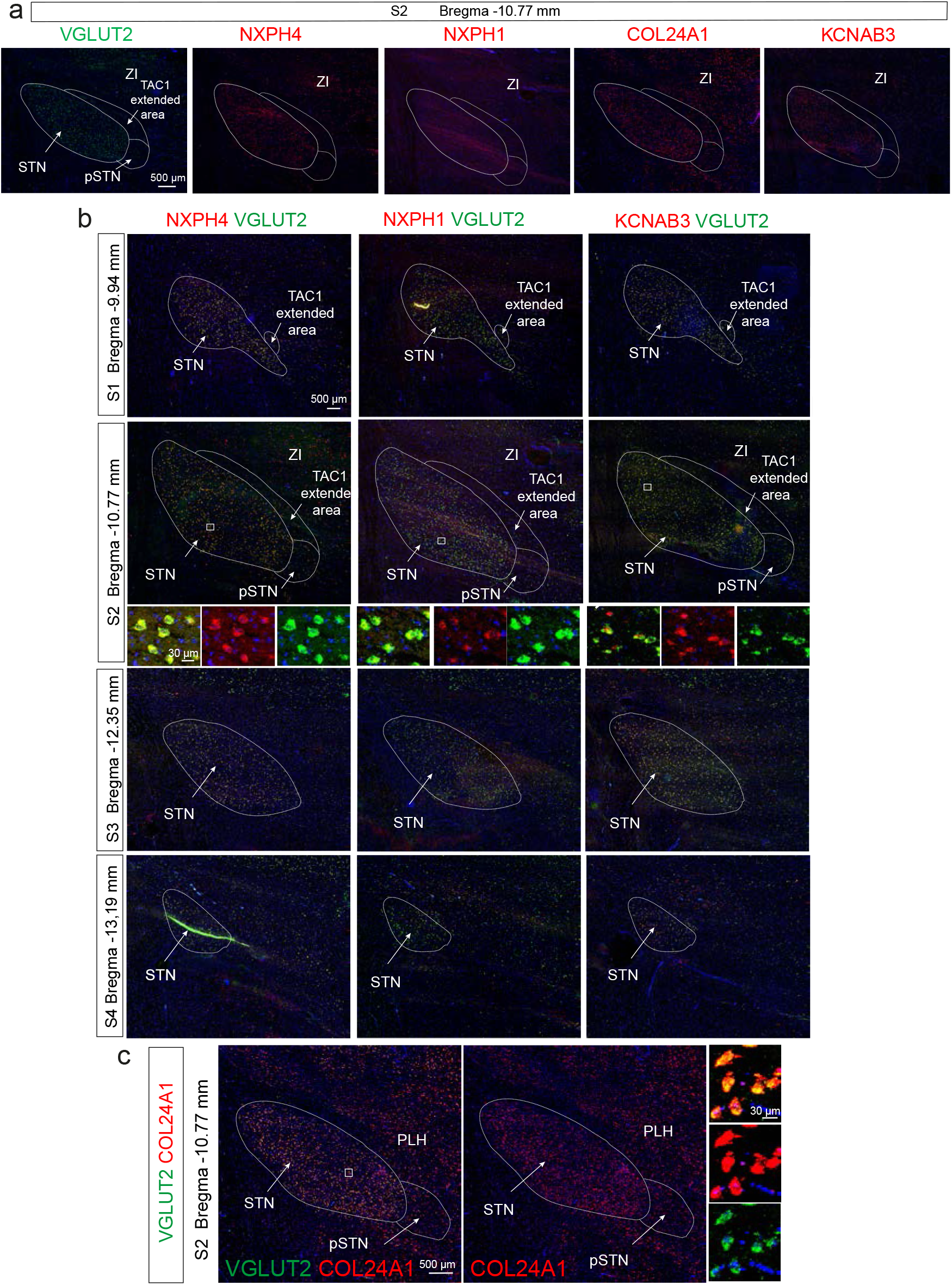
Conservation in macaque of molecular markers originally identified in mouse STN. FISH analysis of molecular markers previously reported as STN-selective in mouse, assessed across the macaque STN - pSTN region. The zona incerta (ZI) is shown as an anatomical reference. Probe colors indicate fluorophores; single labeling is shown in red or green and co-localization in yellow. Section levels S1 - S4 and corresponding Bregma coordinates are indicated. **(a)** *VGLUT2*, *NXPH4*, *NXPH1*, *COL24A1* and *KCNAB3* at S2. *VGLUT2* labels the STN and extends into the canonical pSTN and *TAC1* extended area. *NXPH4*, *NXPH1* and *KCNAB3* show limited or no labeling in the pSTN and *TAC1* extended area, whereas *COL24A1* is detected throughout these regions and extends into the ZI. **(b)** *NXPH4/VGLUT2*, *NXPH1/VGLUT2*, *COL24A1* and *KCNAB3/VGLUT2* across S1 - S4. *NXPH4*, *NXPH1* and *KCNAB3* co-localize with *VGLUT2* in the STN but show limited or no labeling in the pSTN and *TAC1* extended area. Enlarged views at S2 correspond to the regions indicated by squares. **(c)** *VGLUT2/COL24A1* and *COL24A1* at S2, with an enlarged view of the indicated region. Scale bars, µm; individual values are indicated in each panel. Full mRNA names and probe information are provided in **Supplementary Table 2**. Abbreviations: FISH, fluorescent *in situ* hybridization; PLH, posterior lateral hypothalamus; pSTN, para-subthalamic nucleus; STN, subthalamic nucleus; ZI, zona incerta. S1 - S4 refers to section (S) levels corresponding to approximate Bregma level as specified in Fig. 1d and e.

In contrast to the conserved expression of *NXPH4*, *NXPH1* and *KCNAB3*, *COL24A1* mRNA showed a broad distribution across STN, pSTN and adjacent lateral areas in the macaque brain (**Fig. 3c**). This was different to what we previously reported in the mouse, where the corresponding *Col24a1* showed a ventral STN distribution.

### Spatial relationship between mouse STN and pSTN varies across axes

We next examined the mouse STN - pSTN region across four rostrocaudal levels (S1 - S4) using the same marker panel while also adding some newly described markers defined for the pSTN in mice, *Nts* and *Crh*.

At the representative S2 level, *Pitx2*, *Vglut2* and *Htr2c* were broadly distributed throughout the STN, whereas *Pvalb* and *Col24a1* showed complementary dorsoventral distributions (**Fig. 4a-d**). *Pvalb* was enriched dorsally and *Col24a1* ventrally, corresponding to the previously described STNa, STN_b_ and STN_c_ molecular domains.

**Figure 4.**
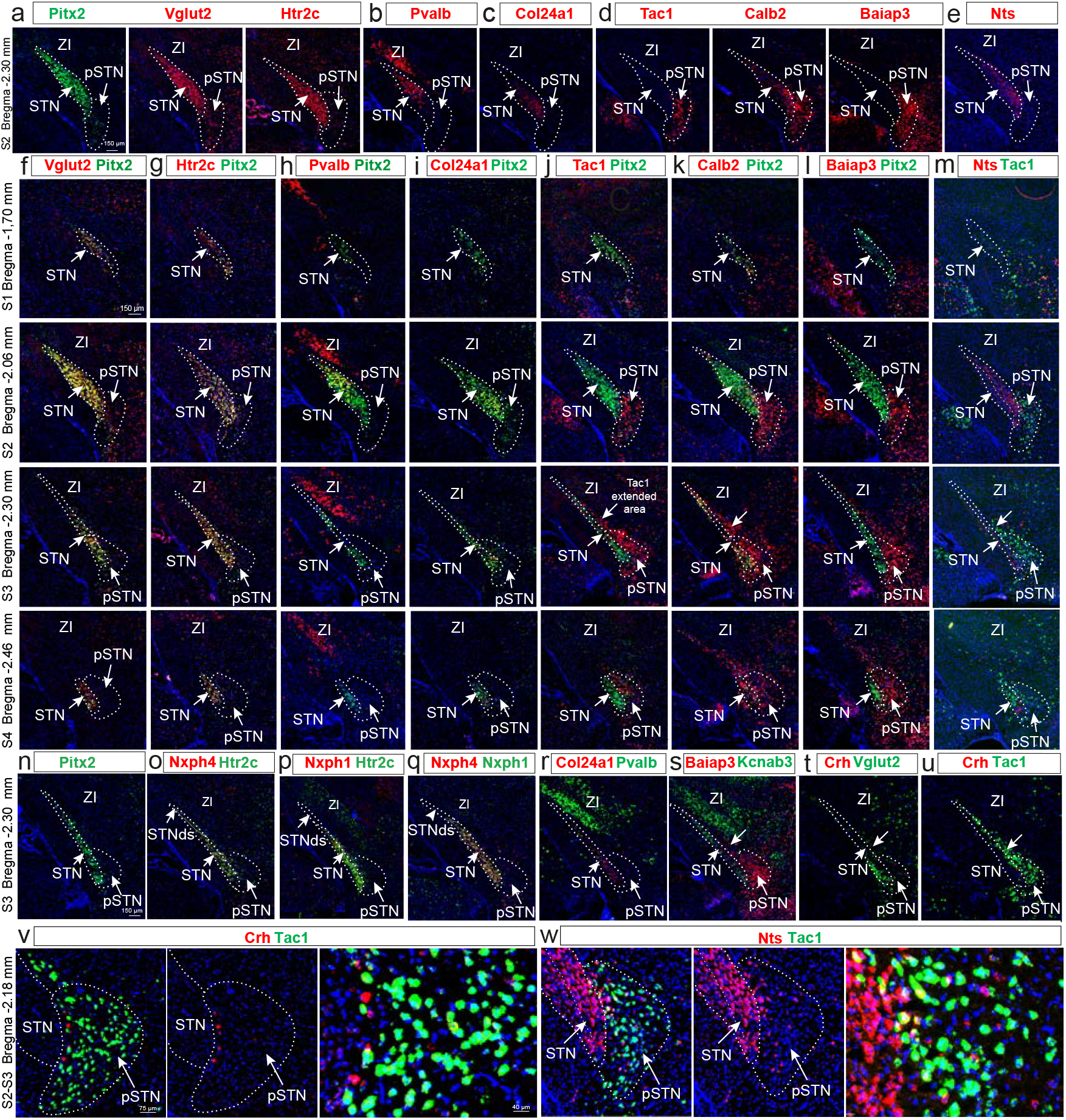
Spatial organization of molecularly defined STN and pSTN domains in mouse. FISH analysis of the mouse STN, canonical pSTN and *Tac1* extended area across section levels S1 - S4. The canonical pSTN and *Tac1* extended area are detected at S2 - S4 but are not detected at S1. The zona incerta (ZI) is shown as an anatomical reference. Probe colors indicate fluorophores; single labeling is shown in red or green and co-localization in yellow. Bregma coordinates and section levels are indicated. **(a)** *Pitx2*, *Vglut2* and *Htr2c*, broadly expressed throughout the STN with reduced labeling in the pSTN. **(b)** *Pvalb*, enriched in the dorsal STN. **(c)** *Col24a1*, enriched in the ventral STN. **(d)** *Tac1*, *Calb2* and *Baiap3*, enriched in the canonical pSTN and extending into the *TAC1* extended area. **(e)** *Nts*, labeling a medial strip across the STN. **(f–m)** FISH across S1 - S4 showing *Vglut2*/*Pitx2* (f), *Htr2c*/*Vglut2* (g), *Pvalb*/*Pitx2* (h), *Col24a1*/*Pitx2* (i), *Tac1/Pitx2* (j), *Calb2/Pitx2* (k), *Baiap3/Pitx2* (l) and *Nts/Tac1* (m). *Pitx2*, *Vglut2* and *Htr2c* are broadly distributed throughout the STN. *Pvalb* and *Col24a1* show complementary dorsal and ventral distributions, respectively. *Tac1*, *Calb2* and *Baiap3* show limited or no co-localization with *Pitx2* and extend dorsally from the canonical pSTN into the *Tac1* extended area. *Nts* labels a medial STN domain and is also detected in the pSTN. **(n–u)** FISH at S3. *Pitx2* (n), *Nxph4/Htr2c* (o), *Nxph1/Htr2c* (p), *Nxph4/Nxph1* (q), *Col24a1/Pvalb* (r), *Baiap3/Kcnab3* (s), *Crh/Vglut2* (t) and *Crh/Tac1* (u). *Nxph1* and *Nxph4* co-localize with each other and with *Htr2c* in STN. *Col24a1* and *Pvalb* show complementary dorsal - ventral distributions with partial overlap in the central STN area. *Baiap3* and *Kcnab3* show limited co-localization: *Baiap3* detected in the pSTN and *Tac1* extended area and *Kcnab3* detected in the STN. *Crh* is detected in rare cells in the pSTN but is not detected in the *Tac1* extended area. **(v,w)** Higher-magnification views of the STN–pSTN border showing *Crh/Tac1* (v) and *Nts/Tac1* (w). *Tac1* extends dorsally beyond the canonical pSTN, whereas *Crh* is absent or rare in this area. *Nts* is abundant in the ventral aspect of STN and less abundant in the pSTN, with *Nts/Tac1* co-localization observed near the STN - pSTN border. Arrows indicate the *Tac1* extended area. Scale bars, µm; individual values are indicated in each panel. Full mRNA names and probe information are provided in **Supplementary Table 2**. Abbreviations: FISH, fluorescent *in situ* hybridization; pSTN, para-subthalamic nucleus; STN, subthalamic nucleus; ZI, zona incerta. S1 - S4 refers to section (S) levels corresponding to approximate Bregma level as specified in Fig. 1d and e.

The distribution of the STN_a_, STN_b_ and STN_c_ domains varied along the rostrocaudal axis. At S1 and S4, the STN_a_ profile predominated; at S2, all three domains were present; at S3, all three remained detectable but the STN structure appeared as a narrow “strip” which was thinner in the medial - lateral dimension than at the S2 level, while equally elongated in the dorsal - ventral dimension (**Fig. 4f–i**).

*Nts* was added to the mouse marker panel based on our original identification of this gene as differentially expressed in the STN and pSTN; in the original snRNA-Seq analysis, *Nts* grouped in the same transcriptomic cluster as the other STN genes analyzed here^33^. Here, we found that *Nts* was present in the anterior extreme of the STN at S1 and posterior extreme at S4, and reached along the full dorsoventral extent of the STN at both S2 and S3. However, at S2, *Nts* was only detected in the medial aspect of the STN while rarely detected in lateral STN, thus forming a medial^high^ – lateral^low^ gradient, appearing as a narrow but elongated strip. At S3, *Nts* mRNA was present in the narrow strip representing the STN structure at this section level (**Fig. 4e, m**). Further, *Nts* mRNA was observed in the canonical pSTN (**Fig. 4e, m**), in accordance with recent report^34^.

*Tac1* delineated the canonical pSTN positioned ventral to the ventral aspect of the *Pitx2*-positive STN (**Fig. 4d**). *Tac1* was not detected at the anterior extreme S1, in accordance with the rostrocaudal distribution of the pSTN in atlases (*see Fig. 1*), but was readily detected from S2 through S4. At S2, *Tac1*-positive cells allowed the identification of the characteristic crescent-shaped pSTN. At S3, in addition to forming the pSTN shape, *Tac1*-positive cells extended as a narrow and elongated population along the STN structure, a narrow “strip” at this posterior section level. At S4, the most posterior level, the *Tac1* area was even more extensive and surrounded the molecularly defined STN, which is small at its posterior extreme (**Fig. 4j**). The narrow *Tac1*-positive area observed at S3 directly bordered the narrow and *Nts*-positive STN “strip” along the dorsoventral axis (**Fig. 4m**).

*Calb2* and *Baiap3* showed a similar distribution as *Tac1* across S1–S4 (**Fig. 4k,l**), but both were less selective for pSTN and extended further into the posterior lateral hypothalamus (**Fig. 4l**).

At S3, the combined territory of the molecularly defined STN (at S3 detected as a narrow “strip” positive for all STN markers described including *Nts*) and the adjacent *Tac1*-positive population approximated the conventional cone-shaped anatomical STN territory observed at the S2 section level, commonly used to depict the STN structure (**Fig. 4j-m, n-p**).

As in macaque, the elongated *Tac1*-positive population aligning the medial STN was designated the “extended *Tac1* area” rather than being classified as an anatomical extension of the pSTN.

Within the canonical pSTN, *Tac1*-positive cells were predominantly *Pitx2*-negative (**Fig. 4l**), and *Tac1*, *Baiap3* and *Calb2* showed partial rather than complete overlap within both the canonical pSTN and the extended *Tac1* area (**Fig. 4k-m, w**). *Crh* was not detected in the extended *Tac1* area (**Fig. 4t**), and *Crh*-positive and *Tac1*-positive cells were largely non-overlapping in the canonical pSTN (**Fig. 4v**).

Finally, whole-brain FISH and light-sheet imaging independently demonstrated the three-dimensional relationship between the *Pitx2*-positive STN and the observed *Tac1*-positive populations. As in our section material, *Pitx2* labeling in the intact mouse brain delineated the continuous STN, whereas *Tac1* labeling revealed the canonical pSTN and an additional population following the medial dorsoventral contour of the STN. As observed in individual sections above, this extended *Tac1* population became more prominent toward posterior levels where it aligned with the medial aspect of the *Pitx2*-positive STN (**Supplementary Videos 1-3**, **Supplementary Figure 5**, frame-by-frame data representation from the whole brain video material).

Corresponding expression patterns for the complete marker set applied in our study were also observed upon querying publicly available images available in the Allen Mouse Brain Atlas ISH and MERFISH datasets (**Supplementary Figure 6**).

### A ventral STN domain is enriched in *HTR2C* mRNA in the macaque

*HTR2C* mRNA showed an apparent species difference in its spatial distribution. In mouse, *Htr2c* mRNA was detected across the STN structure (*see Fig. 4 above*), consistent with our previous observations and public datasets as reported above. In macaque, we found that *HTR2C* mRNA was enriched in the ventral STN across the rostrocaudal axis (**Fig. 5a**).

**Figure 5.**
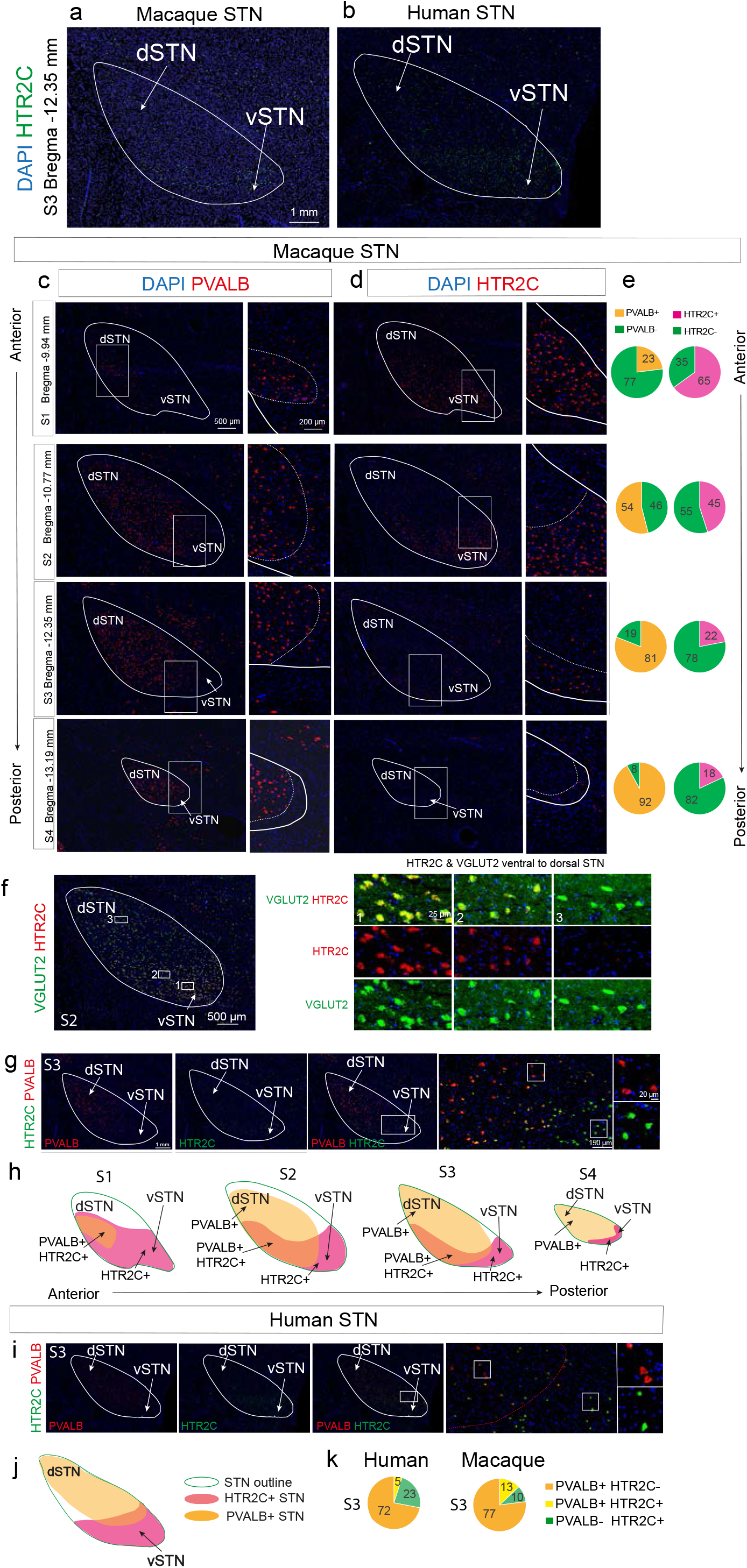
A ventral STN domain is enriched in *HTR2C* in macaque and human. FISH analysis of *HTR2C* and *PVALB* expression in the macaque and human STN. **(a,b)** *HTR2C* expression in macaque (a) and human (b) STN with DAPI counterstaining, detecting more abundant *HTR2C* labeling in the ventral than dorsal STN. **(c,d)** *PVALB* (c) and *HTR2C* (d) expression across macaque section levels S1 - S4. Enlarged views correspond to the indicated regions; dotted lines indicate the boundary between regions with higher and lower labeling. **(e)** Semi-quantitative assessment of manually counted FISH-labeled cells in macaque STN. Percentages are shown relative to the *VGLUT2*-positive cell population. *PVALB*-positive cells are shown in orange, *HTR2C*-positive cells in pink, and *PVALB* and *HTR2C*-negative cells, respectively, in green. Cells from both macaques were counted; however, the distribution shown is based on one male macaque (17 years) because the total number of labeled cells differed between animals and the samples were not pooled. Complete counts are provided in **Supplementary Table 3**. **(f)** *HTR2C/VGLUT2* at S2. Enlarged views show three selected regions spanning the ventral-to-dorsal STN. **(g)** *HTR2C* and *PVALB* at S3, with enlarged views of the ventral STN. Sparse *HTR2C/PVALB* co-labeling is observed in the *HTR2C*- enriched ventral domain. **(h)** Schematic representation of STN domains across S1 - S4 based on spatial anatomy and *PVALB* and *HTR2C* expression; three STN domains identified: *HTR2C*^+^/*PVALB*^-^ (bright pink), *HTR2C*^+^/*PVALB*^+^ (yellow-pink mix), *HTR2C*^-^/*PVALB*^+^ (yellow). **(i,j)** *HTR2C* and *PVALB* expression in human STN at S3 **(i)** and corresponding schematic representation of internal STN domains as shown in h): *HTR2C*^+^/*PVALB*^-^ (bright pink), *HTR2C*^+^/*PVALB*^+^ (yellow-pink mix), *HTR2C*^-^/*PVALB*^+^ (yellow) **(j)**. **(k)** Semi-quantitative assessment of *HTR2C* and *PVALB* expression in macaque and human STN at S3. The distribution patterns are similar between species; statistical comparisons were not performed because of the limited sample size. Complete counts are provided in **Supplementary Table 3**. Scale bars, µm; individual values are indicated in each panel. Abbreviations: FISH, fluorescent *in situ* hybridization; dSTN, dorsal STN: pSTN, para-subthalamic nucleus; STN, subthalamic nucleus; vSTN, ventral STN; ZI, zona incerta. S1 - S4 refers to section (S) levels corresponding to approximate Bregma level as specified in Fig. 1d and e.

*HTR2C* mRNA showed a similar ventral enrichment in human sections containing the STN (**Fig. 5b**), although the limited human material did not permit assessment of the extent or variability of this distribution across the anteroposterior axis.

We next examined *PVALB* and *HTR2C* across anteroposteior axis (S1–S4) in the macaque brain (**Fig. 5c, d**). As described above, *PVALB* mRNA has been reported as a marker for the dorsal STN in the primate. With the observation of a ventral enrichment of *HTR2C*, it was of particular interest to assess their distribution pattern relative each other. We observed near opposite distribution across the STN structure, with *PVALB* dominating dorsally and posteriorly (**Fig. 5c, e**) and *HTR2C* dominating ventrally and anteriorly (**Fig. 5d-e**). *HTR2C* labeling formed a rather distinct border at all section levels (**Fig. 5d-e**). *HTR2C* dominated at the anterior extreme (S1) and *PVALB* at the posterior extreme (S4). *HTR2C* and *PVALB* partly overlapped at S2 and S3, and *HTR2C* was less prominent at posterior levels where *PVALB* dominated. However, *PVALB* was excluded from the most ventral portion also at the posterior extreme (S4) where only *HTR2C* mRNA was deteted of the two markers (**Fig. 5c-e**).

Cell counting verified the visual observation of near opposite mRNA distribution (**Fig. 5e, Supplementary Table 3**).

Both *PVALB*- and *HTR2C*-positive populations showed extensive overlap with *VGLUT2* (**Fig. 5f**) while limited overlap was observed between the two populations (*HTR2C* vs. *PVALB*) (**Fig. 5g**).

Summarizing the observed distribution of *HTR2C* and *PVALB* across S1 - S4 visualizes their molecular distribution into spatio-molecular domains across the macaque STN (**Fig. 5h**).

Limited human material showed the same distribution of *HTR2C* and *PVALB* as observed in macaque (**Fig. 5i, j**). Cell counting in both primate species at the S3 level (the only available section level in the human) confirmed similar distribution patterns (**Fig. 5k**).

Querying the macaque dataset by Allen Brain Cell (ABC) Atlas independently supported our observation of a ventral *HTR2C* mRNA enrichment in the macaque STN (**Supplementary Figure 2**) while this marker was not searchable in the publicly available human pre-print dataset by Allen Brain Cell (ABC) Atlas (https://knowledge.brain-map.org/abcatlas).

### Spatial relationship of *TAC1*-positive populations with MFB

We next examined the spatial relationship between the *VGLUT2* and the *TAC1* labeled areas (the canonical pSTN and the extended *TAC1* area) and the MFB, which traverses the medial aspect of the STN - pSTN region (**Fig. 6a**).

**Figure 6.**
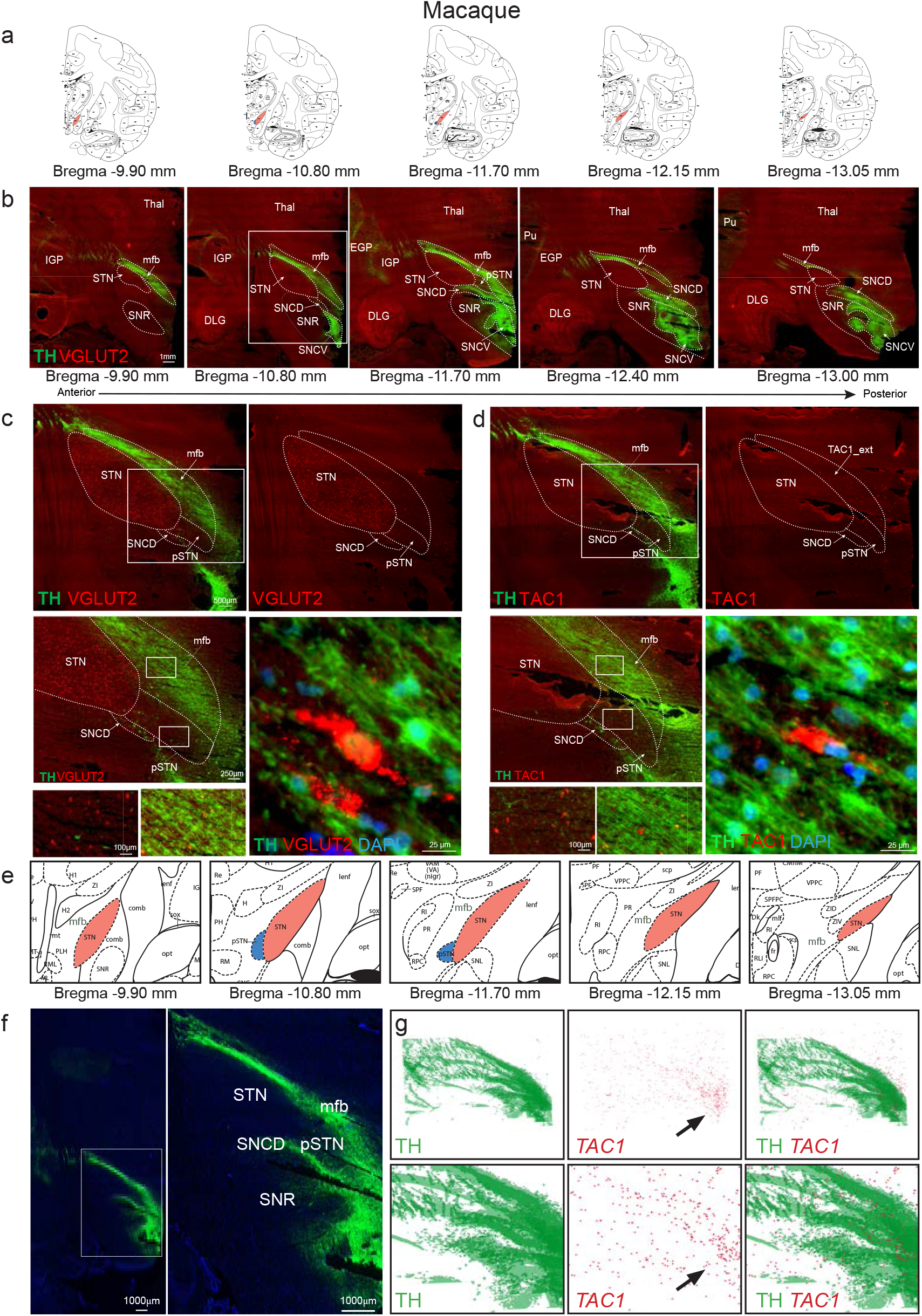
Spatial relationship between *TAC1*-positive cells and the medial forebrain bundle. Combined FISH for *VGLUT2* and *TAC1* and immunofluorescence for tyrosine hydroxylase (TH-IR) to visualize the presence of cell bodies in the TH-positive fibers of the medial forebrain bundle (MFB). Probe colors indicate fluorophores; single labeling is shown in red or green and co-localization in yellow. Bregma coordinates are indicated. **(a)** Macaque atlas sections spanning the anteroposterior axis. Bregma coordinates as indicated in atlas^41^. The medial forebrain bundle (mfb in maps, MFB in text) was manually annotated; STN and pSTN labels were manually adapted from the original atlas terminology. **(b)** *VGLUT2* FISH combined with TH-IR in serial sections. *VGLUT2*-positive cell bodies are detected in the STN and thalamus (Thal), whereas TH-IR identifies fibers and TH-positive cell bodies in the substantia nigra *pars compacta* (SNC), including dorsal and ventral tiers (SNCD and SNCV). At Bregma −10.80 mm, *TAC1*-positive cell bodies are located in close spatial association with TH-IR-positive fiber bundles. **(c)** TH-IR/*VGLUT2* at Bregma −10.80 mm. Enlarged views show *VGLUT2*-positive cell bodies in relation to TH-IR-positive fiber bundles. **(d)** TH-IR/*TAC1* at Bregma −10.80 mm. Enlarged views show TAC1-positive cell bodies in relation to TH-IR-positive fiber bundles. **(e)** Atlas sections at the same Bregma levels as in (a), with enlarged views of the STN - pSTN region^41^ **(f)** TH-IR in a representative section for the serial reconstruction, shown for anatomical orientation. **(g)** Serial reconstruction of TH-IR and TAC1 *signals* in a specimen-specific 3D coordinate system using multiple sections with TH-IR and TAC1 FISH. TH signal is shown as a green field and TAC1 signal as a red point cloud; arrows indicate regions of high TAC1 signal density. The merged reconstruction shows TAC1 signal within and immediately adjacent to the TH-positive fiber territory along the medial aspect of the STN. One analysed section contained both TAC1 and TH measurements, providing direct spatial correspondence between the two signals. Scale bars, µm; individual values are indicated in each panel. Abbreviations: DLG, dorsolateral geniculate nucleus; EGP, globus pallidus externa; FISH, fluorescent *in situ* hybridization; MFB (mfb), IGP, globus palidus interna; medial forebrain bundle; pSTN, para-subthalamic nucleus; Pu, putamen; TH-IR, tyrosine hydroxylase immunoreactivity; SNC, substantia nigra *pars compacta*; SNCD, substantia nigra *pars compacta*, dorsal tier; SNCV, substantia nigra *pars compacta*, ventral tier; SNR, substantia nigra *pars reticulata*; STN, subthalamic nucleus; Thal, thalamus. For additional abbreviations in atlas images, see original publication^41^.

Immunoreactive TH-positive fiber bundles of the MFB were observed along the medial aspect of the STN and canonical pSTN, reaching along the medial aspect of the STN (Fig. 6b). *TAC1*-positive cells were detected within the same brain area; that is, *TAC1* FISH labeling was intermixed with immunoractivity for TH (**Fig. 6c-d**).

Using *VGLUT2* to outline the STN structure, digital three-dimensional analysis of sections containing *TAC1*-positive cells and TH-positive fibers (**Fig. 6e, f**) along the rostrocaudal axis (**Fig. 6e**) was used to further characterize their spatial relationship, showing that *TAC1* labeling was detected within the TH-positive domain (**Fig. 6g**; **Supplementary Figure 7**, **Supplementary Video 4, 5**).

Within this region, the *TAC1*-signal followed the medial contour of the *VGLUT2*-defined STN across successive section levels. TH-positive fibers coursed among *TAC1*-signal, supporting spatial intermingling of the two signals. The three-dimensional reconstruction demonstrates regional spatial correspondence. Together, these observations show that the *TAC1*-positive population occupies a region closely associated with the TH-positive component of the MFB along the medial aspect of the STN.

Whole-brain imaging in the mouse similarly demonstrated *Tac1*-positive cells within the region occupied by the TH-positive fiber system along the medial aspect of the STN, whereas *Pitx2*-positive cells were absent or rare in this region (**Supplementary Figure 5, Supplementary Video 6, 7**). This spatial relationship between *Tac1* and TH was maintained across consecutive anteroposterior frames and was therefore not restricted to a single imaging plane.

### Molecularly supported maps of the subthalamic region

Based on our observations, we summarized the individual gene expression patterns for both species (**Table 1**) and integrated the molecular distributions identified in macaque (**Fig. 7a**) and mouse (**Fig. 7b**) into molecularly enhanced spatial maps of the subthalamic region, paralleled with published atlases for macaque (**Fig. 7a**) ^41^and mouse (**Fig. 7b**) ^42^ for full outline of the anatomical area. Applying the battery of markers studied, the maps depict the molecularly defined STN, canonical pSTN, extended *TAC1* area and their spatial relationship to the TH-positive MFB.

**Figure 7.**
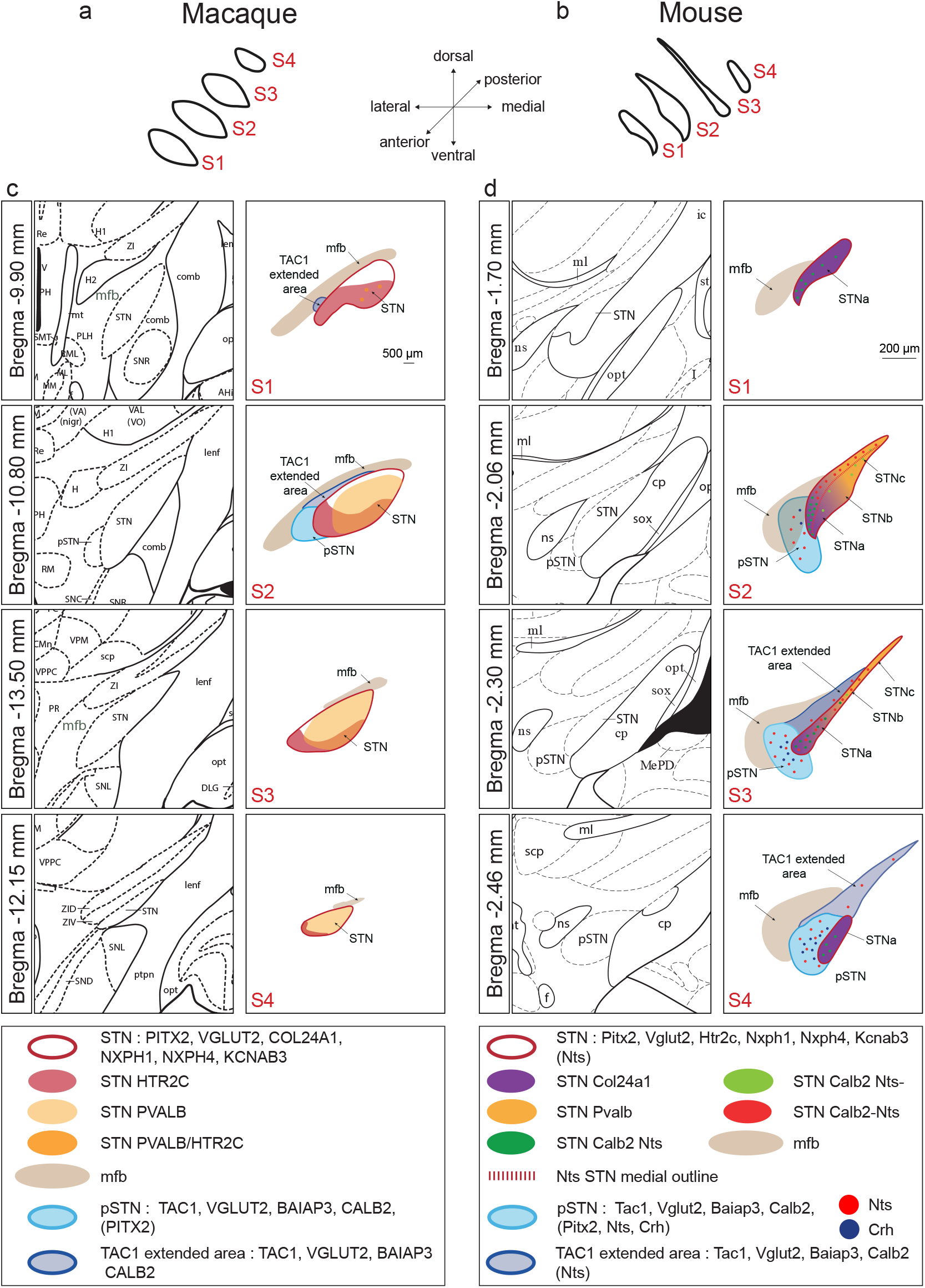
Molecularly supported anatomical maps of the subthalamic area in macaque and mouse. Schematic outlines of the STN across macaque **(a)** and mouse **(b)**, with spatial orientation planes represented in middle. Brain atlas representations (*left*) paralleled with illustrated molecular information provided throughout this study (*right*) for the macaque **(c)** and mouse **(d)** subthalamic region at the indicated Bregma levels. Gene expression patterns identified in this study are based on the atlas representations to delineate the the STN, canonical pSTN and MFB, while molecularly defined internal domains and molecularly identified populations are drawn based on molecular data presented in the study. The maps provide transcriptomic detail that complements the anatomical structures represented in the brain atlases. Color coding is indicated in the figure legend. Full mRNA names are provided in **Supplementary Table 2**. Atlas sources: Macaque^41^, mouse^42^. Abbreviations: pSTN, para-subthalamic nucleus; mfb, medial forebrain bundle; STN, subthalamic nucleus. STN_a_, STN_b_, STN_c_ refer to spatio-molecular domains in mouse STN (original publication^33^). For abbreviations in atlas images, see original publications. S1 - S4 refers to section (S) levels corresponding to approximate Bregma level as specified in Fig. 1d and e.

**Table 1.**
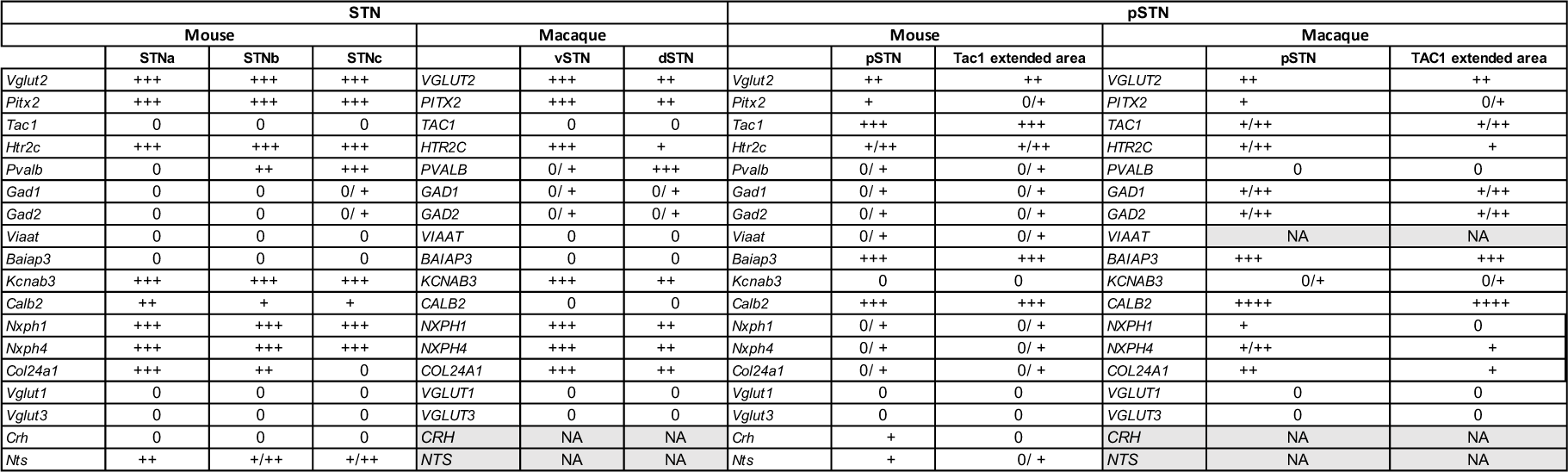
Summary of molecular marker distribution across the STN and pSTN in macaque and mouse. Summary of the distribution of molecular markers across the subthalamic nucleus (STN), para-subthalamic nucleus (pSTN) and molecularly defined internal domains in macaque and mouse. STN_a_, STN_b_ and STN_c_ denote internal spatio-molecular STN domains (^33^), vSTN and dSTN denote ventral and dorsal STN domains, respectively. pSTN denotes the canonical para-subthalamic nucleus as defined in brain atlases. The *TAC1* (*Tac1 in mouse*) extended area denotes a *TAC1*-positive territory identified in both species in the present study that extends beyond the canonical pSTN and expressing several molecular markers associated with the pSTN. FISH signal density is based on visual inspection of cells positive for FISH signal, and indicated as follows: 0, no detectable signal; +, low; ++, moderate; +++, high. Marker distributions are based on the expression patterns observed in the material analysed in this study.

In macaque, *HTR2C* and *PVALB* defined largely complementary molecular territories within the molecularly defined STN, with *HTR2C* mRNA distribution enhanced in a distinct ventral domain the STN. In mouse, the STN showed the previously characterized *Pvalb*/*Col24a1* organization across the dorsoventral axis, and the herein described medially located narrow “strip” of *Nts* mRNA; these differential expression patterns displayed as gradients in the dorsoventral and mediolateral planes, respectively. In both species, the extended *TAC1* area followed the medial contour of the molecularly defined STN and occupied a region containing TH-positive fibers, a finding validated in 3D reconstructions. The anatomical identity of this extended *TAC1*-positive region warrants further investigation to determine whether it represents a distinct anatomical area, an extension of the pSTN, or a broader component of the posterior lateral hypothalamus (**Supplementary Figure 8**, presenting these alternatives in a simple model).

Expression patterns accessible in the represented Allen Institute datasets were concordant with the corresponding experimental observations (**Supplementary Figures 2, 3, 6**). Note that not all mRNAs were searchable in the macaque and human datasets, and that pSTN is not an annotated structure in the Allen Institute datasets; thus the supplementary figures only present the STN structure as the annotated structure.

## Discussion

Our comparative molecular mapping supports an emerging view of the STN as a spatially organized and molecularly heterogeneous structure in which a largely conserved molecular identity coexists with species-specific specialization (**Fig. 7**). Across mouse and macaque, the molecularly defined STN was characterized by predominant *VGLUT2* and *PITX2* expression, together with broadly distributed STN-associated markers including *NXPH1*, *NXPH4* and *KCNAB3*. Against this conserved framework, *HTR2C* showed an unexpected difference in spatial organization.

Whereas *Htr2c* was broadly distributed across the mouse STN, *HTR2C* was enriched in the anterior and ventral macaque STN, with a similar localized *HTR2C* mRNA pattern detected in the human STN sample examined. Further, while not addressed in detail, querying images published in the ABC Atlas provided support for our detections. Importantly, the principal species difference identified here was not the presence of *HTR2C* mRNA in the STN, which has been long described, but its localized spatial distribution within the STN structure.

This finding adds a molecular dimension to the longstanding question of how the primate STN is organized. Anatomical and connectivity studies have described sensorimotor, associative and limbic territories distributed broadly along the dorsolateral to ventromedial axis, but whether these represent discrete subdivisions or overlapping spatial gradients remains unresolved^12,17,43–45^. Our identification of a ventral STN domain enriched in *HTR2C* mRNA and forming largely complementary patterns to the dorsal *PVALB*-enriched population provides an independent molecular feature with which these organizational models can be further investigated. While limited overlap was observed between these two markers, both markers were indeed present in cells of the central aspect of the STN structure. Thus, *PVALB* and *HTR2C* mRNAs allowed the tentative identification of three molecularly supported anatomical domains within the macaque and human STN: The *HTR2C*^+^ domain, most prominent at the ventral STN, the central *HTR2C*^+^/*PVALB*^+^ domain, and the *PVALB*^+^ domain, most prominent dorsally. In summary, these three spatio-molecular domains of the macaque STN were based on gene expression gradients and their relative distribution varied substantially across the axes (**Fig. 7**, **Table 1**).

*HTR2C* and *PVALB* positive cells were predominantly *VGLUT2*-positive, indicating molecularly distinct populations within the glutamatergic STN rather than fundamentally different transmitter phenotypes. *VGLUT2* was consistently detected as more dense in ventral than dorsal STN, in agreement with a previous report^11^. Of further importance, molecular composition varied not only along the dorsoventral axis but also along the rostrocaudal axis, with *HTR2C* expression predominating in the rostral (anterior) STN and *PVALB* expression increasing toward the caudal (posterior) STN. At the anterior extreme of the STN, *HTR2C* was by far most dominant of the two, reaching across almost the full extent of the STN at this level.

Importantly, while providing localized mRNA patterns within the STN structure, our data do not establish that the *HTR2C*-enriched population corresponds to an associative or limbic territory, but identify a molecular landmark with which such relationships can be investigated.

*HTR2C* expression in the STN itself is not novel. *Htr2c* expression and *5-HT2C* receptor binding have been demonstrated in the rodent STN, and *HTR2C* mRNA has been mapped more broadly in the primate basal ganglia^20,46–50^. These studies did not, however, identify a spatially restricted *HTR2C* population within the primate STN. Our findings therefore extend previous observations by identifying a prominent anterior/ventral *HTR2C*-enriched domain in the STN of the macaque, with a similar distribution in the limited human tissue examined. This is a molecular anatomical finding and does not establish receptor protein abundance, serotonergic innervation, receptor function, or a specific behavioral role. The identification of this spatially restricted population nevertheless provides a defined molecular feature for future investigation of *5-HT2C* signaling within the primate STN.

Spatio-molecular gradients within the STN structure were also considerable in the mouse. The relative distribution of STN_a_, STN_b_, STN_c_ domains, previously described based on oppsite expression gradients along the dorsoventral axis of *Col24a1* (ventral^high^) and *Pvalb* (dorsal^high^)^33^, changed substantially between anterior and posterior levels, such that the molecularly defined STN could not be represented adequately by a single transverse organization. In addition, a new type of internal STN gradient was observed when we now introduced *Nts* (neurotensin) in the FISH analysis. We originally reported *Nts* in two of six clusters detected as differentially expressed in the *Pitx2*-targeted snRNA-Seq experiment^33^. Notably, *Nts* was detected in both the STN-enriched *Pitx2*-positive red cluster (which identified several STN marker genes, including *Col24a1*, *Htr2c*, *Kcnab3*, *Nxph1*, *Nxph4*), and a separate cluster exhibiting low or absent expression of these STN-associated genes. Here, our FISH experiment identified a gradient across the mediolateral planes of the mouse STN, distinguishing a medial STN domain defined by *Nts* (observed as narrow strip along the full dorsoventral axis of the STN). This medial *Nts* expression was detected at the S2 level, where the *Nts* gradient roughly divided the STN into a medial (*Nts^+^*) and lateral (*Nts^-^*) domain. At the more posterior level S3, the full extent of the STN (at this level appearing as a narrow strip) was *Nts^+^*. This mediolateral molecular gradient within the STN was unexpected. For independent validation, we compared all our gene expression patterns with images available in Allen Institute’s ISH and MERFISH repositories; by consulting these data sets, we found that the mRNA patterns we observed in our own material seemed to show the same distribution patterns in these repositories, including the medial distribution of *Nts* mRNA. Together, the findings observed in our own material and in the publicly available dataset reveal that STN molecular identity is shaped by intersecting gene-expression gradients across the rostrocaudal, mediolateral, and dorsoventral axes, highlighting a spatially heterogeneous organization that changes along the whole extent of the nucleus (**Fig. 7**, **Table 1**).

A related picture emerges from the neighboring pSTN. Our previous transcriptomic and FISH analyses identified *Tac1* and other markers distinguishing the pSTN from the molecularly defined STN in mice, and we subsequently showed that *TAC1* also marks the macaque pSTN^32,33^. In literature, *Tac1* in the rodent has long been described as a highly selective marker for the pSTN^35^. Here, *Tac1*, *Baiap3* and *Calb2* identified the molecularly defined canonical pSTN in both species, but of these, only *Tac1* was selective for pSTN and not detected in either STN and the posterior lateral hypothalamus. *Baiap3* and *Calb2* were equally excluded from STN but extended in the posterior lateral hypothalamus in both species. In the mouse pSTN, *Nts* and *Crh* allow additional molecular heterogeneity in the mouse (Shah, Dunning et al. 2022).

These were analyzed here in the mouse but not examined in the primate because of limited material. Our findings are consistent with recent work distinguishing *Tac1*- and *Crh*-expressing glutamatergic populations within the mouse pSTN, with distinct projection and functional properties^40^. Our previous studies further demonstrated distinct projection patterns and behavioral effects of STN and pSTN populations, with STN populations projecting predominantly to basal ganglia structures and the *Tac1*-defined pSTN population showing stronger projections to limbic structures, although both populations had projections to the ventral pallidum and substantia nigra *pars compacta*^32^. Thus, the pSTN is increasingly characterized as a molecularly heterogeneous structure rather than a single uniform population, consistent with its involvement in several different functions, acting as a hypothalamic hub for interoceptive, autonomic and motivated behaviors^34^.

The *TAC1*-positive population was found to extend beyond the canonical pSTN, reaching up along the medial aspect of the molecularly defined STN structure in both mouse and macaque, with the visual appearance of a narrow *TAC1*-positive strip of cells. Thus, *TAC1* expression defines a spatially continuous molecular population that includes the canonical pSTN but also extends beyond its currently defined anatomical boundaries. We refer to this as an extended *TAC1* area rather than assuming that the entire population represents an anatomical extension of the pSTN (**Fig. 7**, **Table 1**). The anatomical identity of this extended territory remains unresolved. Its additional molecular profile positive for *BAIAP3* and *CALB2* in both species is consistent with a hypothalamic identity, but does not establish whether it belongs to the pSTN or represents a molecularly continuous population associated with the broader posterior lateral hypothalamus. In the mouse, while *Tac1* extends along the STN border, *Crh* is does not. Because *Tac1* and *Crh* identify distinct pSTN populations in the mouse, the extended *Tac1*-positive territory could represent an extension of the *Tac1*^+^/*Crh*^-^ pSTN population, or alternatively a distinct population associated with the posterior lateral hypothalamus (**Supplementary Figure 8**, a simple model presented of possible anatomical identity of the *TAC1*-labeled cells).

Resolving these possibilities will require additional molecular and anatomical markers that distinguish the extended territory from the canonical pSTN. Our findings nevertheless provide molecular information relevant to recent discussions of the anatomical relationship between the posterior lateral hypothalamus and medial subthalamic region^51^.

Three-dimensional imaging further clarified the spatial relationship of the *TAC1*-positive population, including the canonical pSTN and the extended *TAC1* area, to the TH-positive fibers of the MFB, which traverse the same medial subthalamic region. In both macaque and mouse, the *TAC1*-positive population occupied the vicinity of TH-positive MFB fibers. These observations establish spatial proximity but do not demonstrate synaptic connectivity, cellular colocalization, or functional interaction. The three-dimensional reconstructions therefore provide an anatomical framework for future studies of how this molecularly continuous *TAC1*-positive population and longitudinal fiber systems are organized within the medial subthalamic region. Importantly, molecular continuity should not be taken to imply anatomical identity: the present data establish that *TAC1*-positive cells extend beyond the canonical pSTN, but do not establish that whether the *TAC1*-positive cells associated with the MFB constitutes part of the pSTN. Thus, the association of *TAC1*-positive cells with the MFB needs further investigation to clarify their anatomical and functional identity.

Observations of images available in Mouse Allen ISH and MERFISH datasets supported our STN- and pSTN-associated expression patterns^52,53^, while the ABC Atlas macaque data supported the ventral *HTR2C* distribution in the STN^54^. These publicly available spatial datasets can be considered alongside recent single-cell and single-cell multi-omic atlases that provide complementary molecular information across species, including the transcriptomic characterization of cell types across the adult human brain ^55^ and the single-cell multi-omic atlas of the adult rhesus macaque brain^56^. The datasets provide useful independent molecular references and allow spatial expression patterns to be examined across species. However, as for all tissue analysis, differences in tissue preparation, spatial resolution, transcript detection, sampling and anatomical annotation limit direct comparison between cohorts. In particular, the pSTN is not provided as an independent anatomical annotation in the publicly available macaque and human datasets considered here. We therefore regard these resources as complementary support for our identified molecular patterns rather than as definitive validation.

Several limitations should be considered. First, the number of macaque brains and human sections was limited, and the human analysis was based on tissue from a single individual. The primate and human findings should therefore be considered descriptive and hypothesis-generating, particularly with respect to inter-individual variability. In addition, the mouse brains were derived from young cohorts while the primate material was older age; thus the age factor may play a role in differences observed across species (*Htr2c* vs. *HTR2C*, *Col24a1* vs. *COL24A1*). Future studies will need to address comparable ages. Second, mRNA expression does not establish protein abundance, receptor activity, neuronal connectivity or functional significance. This is particularly relevant for *HTR2C*, for which our data establish a spatial expression pattern but not receptor protein abundance or receptor-mediated serotonergic signaling. Similarly, spatial proximity between *TAC1*-positive cells and TH-positive MFB fibers does not demonstrate anatomical or functional connectivity.

Third, although publicly available spatial transcriptomic datasets provided independent support for those expression patterns that were searchable, differences in species, tissue preparation, sampling and anatomical annotation limit their use as definitive validation. Finally, the molecular boundaries identified here do not necessarily correspond to functional or connectivity-defined territories, and the identity and functional significance of the extended *TAC1* area remain to be established. *Nts* and *Crh* were examined only in mouse because of limited primate material and therefore remain to be investigated in the macaque and human subthalamic region.

The anatomical and molecular heterogeneity identified here may ultimately be relevant to neuromodulation, but any clinical implications remain speculative. The STN and its neighboring regions form a spatially complex environment in which molecularly distinct neuronal populations and fiber pathways occur in close proximity, a feature relevant to understanding the cellular and circuit specificity of neuromodulatory interventions such as DBS^51,57^. Clinical studies indicate that the effects of STN-DBS on non-motor symptoms can vary with stimulation location and network engagement^8^, while connectomic studies further emphasize the contribution of distributed circuits to clinical effects^58,59^. This may be particularly relevant to affective symptoms such as depression in PD, for which outcomes following STN-DBS are variable^60,61^. The proximity of the extended *TAC1* population to the MFB is also of interest given the investigation of MFB stimulation for treatment-resistant depression^62^. However, our study does not establish functional relationships and clinical relevance. Rather, these molecular and anatomical landmarks provide a framework for future studies examining how distinct neuronal populations and fiber pathways contribute to the effects of subthalamic neuromodulation.

In summary, comparative molecular mapping identifies a largely conserved molecular framework of the STN across mouse and macaque while revealing substantial spatial specialization and molecular heterogeneity within the STN and its surrounding subthalamic region. In the primate STN, complementary *HTR2C*- and *PVALB*-enriched expression patterns define distinct molecular territories, with a prominent ventral *HTR2C*-enriched domain also observed in the limited human tissue examined. In the mouse, molecular organization was similarly spatially complex, with previously defined dorsoventral gradients varying along the rostrocaudal axis and a distinct mediolateral gradient revealed by *Nts* expression. Together, these findings indicate that STN molecular identity is shaped by intersecting spatial gradients rather than a uniform transverse organization. The neighboring pSTN was likewise molecularly heterogeneous, with *TAC1*-, *BAIAP3*-, and *CALB2*-associated populations defining the canonical pSTN in both species while cells positive for these markers extended continuously beyond the canonical pSTN along the medial aspect of the STN. Three-dimensional imaging further resolved the spatial relationship of the *TAC1*-positive population (canonical pSTN and/or the extended area) to the TH-positive MFB. Together, these findings provide a molecular and spatial framework, illustrated in molecularly enhanced anatomical maps side by side with published atlas images at same level (**Fig. 7)**. This framework should be useful for better understanding of the three-dimensional organization of the STN and surrounding subthalamic region and establish molecular landmarks for future investigation of their connectivity and circuit organization across species.

## Methods

### Mouse handling and mouse brain tissue samples

All experimental procedures using mice followed Swedish Legislation (Animal Welfare Act SFS 1998:56) and European Union Legislation (Convention ETS 123 and Directive 2010/63/EU). 16 wildtype mice (C57BL/6NTac,Taconic) were used for this study; 8 male and 8 female mice, all at the age of postnatal day 28 (P28). Brains were quickly dissected from mice euthanized by cervical dislocation, snap-frozen in cold (−30°C to −35°C) 2-methylbutane (R99%, Honeywell) and stored at - 80°C until usage. Coronal serial sections were prepared on a Leica CM1950 Cryostat at 16 μm thickness and placed onto Superfrost glass slides in series of 8 slides (Thermo Fisher). The whole STN Bregma (mm) −1.70 to −2.45 was sectioned in each mouse. 16 mouse brains generated 95 slides; 6 sections per slide. In total, N=570 STN sections were analysed. The MFB above the STN and the whole pSTN are included within these sections. Section levels (S) S1 - S4, referring to approximate Bregma (mm): −1.70 (S1), −2.05 (S2), −2.30 (S3), −2.46 (S4) were selected for illustrations.

### Macaque brain tissue samples

Non-human primate brain tissue samples were obtained from two brains derived from adult macaque monkey, *Macaca mulatta*. One male macaque monkey, age 10 years, and one female macaque monkey, age 17 years. No animal was sacrificed for the present study, instead only left-over brain material from a previous study was used.

Animal care was carried out in strict accordance with the European Union Directive of 2010 (Council Directive 2010/63/EU) for care and use of laboratory animals. The authorisation for conducting our experiments was approved by the Ethics Committee in Animal Experiment Charles Darwin (agreement no. Ce5/2011/014).

The protocol was designed to minimize the number of animals and reduce their suffering and distress. The animals were tranquilised with ketamine 500 (10 mg/kg), received morphine (dolorex, 0.04 mg/kg) and were then deeply anesthetized with isoflurane (3%) and oxygen (50%). Brain was removed from the skull and divided into several blocks. Small blocks containing the entire extent of the STN (cube with 18 mm sides) were removed from the skull and cut on a cryostat at 16 µm (**Supplementary Figure 1**).

Sections were mounted on slides and stored at −80°C. Sections including the whole STN (Bregma −9.45; Bregma −13.50) were collected, one section per slide. N=150 sections from the male macaque and N=80 sections from the female macaque were analysed. The MFB above the STN and the whole pSTN are included within these sections. Section levels (S) S1 - S4, referring to approximate Bregma (mm): −9.74 to - 10.13 with a mean value of −9.94 (S1); −10.37 to −11.17 with a mean value of - 10.77 (S2); −12.13 to −12.57 with a mean value of −12.35 (S3); −13.04 to −13.33 with a mean value of −13.19 (S4). Approximate Bregma for each figure is shown in each figure and summarized in **Supplementary Table 1**.

### Human brain tissue samples

Post-mortem human brain sections encompassing the STN (N=3 sections) were obtained from one individual brain collected in a Brain Donation Program of the Brain Bank “GIE NeuroCEB” (GIE Neuro-CEB BB-0033-00011) run by a consortium of Patients Associations: ARSEP (Association for research on multiple sclerosis), CSC (Cerebellar ataxias), France Alzheimer, and France Parkinson. Consent was signed by patient or next of kin in accordance with the French Bioethical Laws. The Brain Bank GIE NeuroCEB has been declared at the Ministry of Higher Education and Research and has received approval to distribute samples (agreement AC-2013-1887), France. ID: 7582, 74 y, post-mortem interval 24 h. This individual had no reported neurological and psychiatric disease.

### Probe and primer information for fluorescent *in situ* hybridization (FISH) on brain sections

Full names of mRNAs and their abbreviations, corresponding probe sequence and primer information for probe generation, see **Supplementary Table 2**.

In the main text, primate mRNA nomenclature (capital letters, italics) is used primarily to facilitate readability when both primate and mouse species are discussed within the same section.

### FISH experimental procedure on brain sections

FISH protocol: https://dx.doi.org/10.17504/protocols.io.bp2l6y631vqe/v2

Preparation of solutions: https://dx.doi.org/10.17504/protocols.io.dm6gpqko1lzp/v1

Briefly, cryo-sections were air-dried and fixed in 4% paraformaldehyde, after which they were acetylated in 0.25% acetic anhydride/100mM triethanolamine (pH 8) and washed in PBS. Sections were hybridized for 18 h at 65°C in 100 μl of hybridization formamide-buffer containing 1 μg/ml digoxigenin (DIG) labeled and 1 μg/ml fluorescein labeled probes for detection of mRNA. Sections were washed at 65°C with SSC buffers (5x and 0.2x), followed by blocking with 20% heat-inactivated fetal bovine serum and 1% blocking solution (Roche #11096176001) in maleic acid buffer. Sections were next incubated with horse radish peroxidase (HRP)-conjugated anti-DIG Fab fragments (Sigma-Aldrich #11207733910) at a dilution of 1:2500 in the same blocking buffer). Signal detection was obtained using Cy3-tyramide at a dilution of 1:100. Fluorescein epitopes were then detected with HRP conjugated anti-fluorescein Fab fragments (Sigma-Aldrich #11426346910) at a dilution of 1:5000 in PBST and revealed using Cy2-tyramide at a dilution of 1:250. Cy2-tyramide and Cy3-tyramide were synthesized as previously described. DAPI was used for nuclear staining.

### Histology and immunofluorescence on brain sections

Selected sections were stained with cresyl violet (0.5%, C0775, Merck Millipore, MA, USA) for histological analysis, or analyzed by immunofluorescence post-FISH.

Tyrosine hydroxylase (TH) immunoreactivity was assessed using anti*-*TH antibody (MAB318, diluted 1:250, Merck Millipore, MA, USA) in FISH blocking buffer overnight at RT. After washes in maleic buffer, 0.1% MABT and PBS, sections were incubated with biotinylated goat anti-mouse IgG antibody (Vector Laboratories #BP-9200) diluted 1:400 in PBS, 1 h at RT. Sections were next washed in PBS and incubated in streptavidin-HRP (Vector Laboratories #SA-5004) in PBS for 1 hr. Revelation of signals was performed using Cy2-tyramide at a dilution of 1:250.

### Slide scanning and analysis

Slides were scanned at 20x magnification on a NanoZoomer 2.0-HT (Hamamatsu), with a resolution of 456 nm/px (55701 DPI). The Ndp2.view software (Hamamatsu) was employed for viewing the images. Published atlases were used for guidance of anatomical structures, as referenced below.

### Manual counting of FISH-labeled cells in brain sections

Manual cell counting of FISH-labeled cells were performed using the NDP.view2 and QuPath softwares based on visual inspection of FISH positive signals. FISH signal was visually observed and rated as: 0, no detectable signal; +, low; ++, moderate; +++, high. Protocol for manual cell counting of FISH-labeled cells: https://dx.doi.org/10.17504/protocols.io.e6nvwqy87vmk/v3

### Whole mouse brain *in situ* hybridization and 3D imaging by lightsheet microscopy

Whole mouse brain *in situ* hybridization was performed by Gubra according to previously published protocols^63^. HCR™ RNA probes targeting mRNA encoded by *Tac1* (Gene ID: 21333), *Pitx2* (Gene ID: 18741), *Vglut2* (Gene ID: 140919) were designed to target all splice variants using a custom probe design proprietary to Molecular Instruments. Brains were imaged on a Luxendo LCS-SPIM (Bruker) light-sheet microscope at 4.4x magnification with 10 µm z-stack intervals. Imaris (Bitplane) software was used for 3D rendering and for generating movies and images. *Tac1*, *Pitx2*, and *Vglut2* signals were mapped to the LSFM reference brain atlas ^64^ together with data from TH whole brain immunolabeling previously generated^65^.

### Anatomical nomenclature and atlas mapping

Anatomical regions and fiber tracts were identified and mapped with reference to the Allen Mouse Brain Reference Atlas ^52^ and the stereotaxic atlases of Paxinos and colleagues: *The Mouse Brain in Stereotaxic Coordinates* ^42,66^ and *The Rhesus Monkey Brain in Stereotaxic Coordinates* ^41^. The canonical anatomical location of the pSTN, which is not represented in all atlases, was identified with reference to the corresponding Paxinos atlas for each species.

For consistency with standard neuroanatomical usage and readability, fiber tracts, pathways, and projections are presented using uppercase acronyms in the main text and figures, while the underlying atlas nomenclature was retained when referring specifically to atlas annotations (e.g., MFB for medial forebrain bundle in main text; mfb in illustrations).

### Publicly available gene-expression datasets

Gene expression patterns observed in our experiments were compared with publicly available gene-expression and spatial transcriptomic datasets from the Allen Institute. Mouse *in situ* hybridization (ISH) gene-expression data were obtained from the Allen Mouse Brain Atlas^52^. Genes of interest were queried using their gene symbols, and the corresponding ISH experiments were reviewed. For each gene, coronal sections were selected, and relevant ISH images were downloaded from the image viewer for figure preparation.

Spatial transcriptomic data for mouse, macaque, and human were queried using the Allen Brain Cell (ABC) Atlas. For mouse, the MERFISH-C57BL6J-638850 dataset with imputed genes and reconstructed coordinates was used^53^. For macaque and human, gene expression patterns were queried in the Human and Mammalian Brain Atlas (HMBA) basal ganglia spatial transcriptomic datasets through the ABC Atlas interface^54^. Genes of interest were queried using the “Genes” panel, and spatial expression patterns were visualized using the “Color by” function.

For macaque and human, the subthalamic nucleus (STN) was selected under “Manual anatomical annotations” (labelled “Sth” in the database) to visualize the anatomical boundaries of the STN. The para-subthalamic nucleus (or its abrevation, pSTN) was not available as a separate anatomical annotation and was therefore not independently selected. Resulting images were exported as PNG files for figure preparation.

### Data analysis and brain rendering

Three-dimensional representations of the STN and para-subthalamic nucleus (pSTN; designated PSTN or PSTh in the source atlases) were generated separately for mouse and macaque monkey.

### Atlas sources and anatomical definitions

Mouse structures were obtained directly from the Allen Mouse Brain Common Coordinate Framework annotation volume using structure identifiers 470 and 364, respectively. A 50-µm isotropic annotation volume was used for the final multiview rendering series. Binary masks were generated independently for both structures by retaining voxels that matched the corresponding structure identifier. A triangular surface was extracted from each binary mask with the marching-cubes algorithm at an isovalue of 0.5. Vertex coordinates were transformed from voxel indices into atlas space using the affine transformation stored with the annotation volume. The resulting surfaces were smoothed using a windowed-sinc surface-smoothing operation implemented through VTK/Vedo. Fourteen smoothing iterations were used to reduce voxel stair-stepping while retaining the atlas-derived topology and relative position of the two nuclei.

The macaque reconstruction was based on serial coronal boundaries in the Paxinos macaque stereotaxic atlas because the pSTN is not provided as a separate region in the NIMH Macaque Template, the Subcortical Atlas of the Rhesus Macaque, or the D99 atlas. For the macaque monkey, the boundaries labelled STh and PSTh were extracted from the supplied Paxinos coronal atlas plates. The STN was traced on atlas figures 63–72, whereas the pSTN was identifiable on figures 66–68.

Consecutive plates were separated by approximately 0.45 mm in the anteroposterior direction. Each identified region was converted into a binary two-dimensional mask while preserving the original atlas boundary. Coordinates in each plate were converted from page coordinates into anatomical millimetres using the printed stereotaxic grid. The midline was located at page coordinate 305.78 points, the dorsoventral zero coordinate at 823.16 points, and the calibration was 18 page points per millimetre. The mediolateral and dorsoventral coordinates were obtained from this calibration, and the anteroposterior coordinate was assigned from the Bregma value of the corresponding atlas plate.

A contour-loft procedure was used to construct continuous three-dimensional surfaces from the serial atlas sections. For each structure, the right-hemisphere boundary was extracted from every available plate and resampled by arc length to 192 equally spaced points. Contour orientation was standardized, and the starting index of every contour was cyclically shifted to minimize the mean squared distance to the preceding contour. This correspondence step reduced twisting of the triangulated surface between adjacent sections. The trajectory of every corresponding boundary point was interpolated along the anteroposterior axis using a shape-preserving piecewise cubic Hermite interpolating polynomial. Intermediate contour rings were generated at 0.05-mm intervals. Adjacent rings were then connected with triangular faces to produce a closed surface. Because the atlas displayed each nucleus on a finite number of plates, terminal contours were added to close the rostral and caudal ends of the reconstructed surfaces. The centre of each terminal contour was estimated by linearly extending the displacement between the two nearest observed contour centres. At the terminal coordinate, the boundary radius was reduced to 2.5% of the nearest observed contour radius, producing a tapered rather than planar closure. The reconstructed anteroposterior extents were approximately −9.00 to −13.95 mm for the STN and −10.35 to −12.15 mm for the pSTN. The meshes were cleaned and smoothed for 18 iterations with a pass-band value of 0.1. Smoothing was restricted to removal of high-frequency surface irregularities and did not introduce additional anatomical landmarks.

### Rendering and camera standardization

Surface visualization was performed using Vedo and the VTK rendering backend. Images were generated with orthographic projection to avoid perspective-dependent changes in apparent size. The STN and pSTN were rendered with plastic surface lighting on a uniform light background. For the rotational series, the camera was positioned initially in a straight posterior view and rotated around the dorsoventral axis in 1° increments through 360° degrees. Dorsal was maintained at the top and ventral at the bottom in every frame. Identical colours and comparable projection settings were used across species; anatomical size was not normalized between species.

### Interpretive limitations

The mouse models are direct surface representations of discrete Allen CCF labels and inherit the anatomical definitions and spatial resolution of that annotation. The macaque models are serial-section contour reconstructions. Their boundaries coincide with the Paxinos outlines at the sampled coronal levels, whereas their geometry between sections was estimated by shape-preserving interpolation. The tapered terminal portions are extrapolations added to obtain closed surfaces^67–69^.

### Light-sheet image extraction and registration

Light-sheet microscopy datasets (supplied by Gubra) contained multiresolution HDF5 (.h5) and Imaris (.ims) files. The high-resolution dataset contained separate channels for tissue autofluorescence, *Tac1*, *Pitx2* and *Vglut2*. An additional dataset contained tyrosine hydroxylase (TH) immunoflurescence from a separate specimen. Channels were extracted individually using Python with h5py, tifffile, NumPy and SciPy. For coronal visualization, images were generated from the native-resolution volumes at an in-plane sampling of 5 × 5 µm. Five adjacent optical planes, covering 25 µm in the anteroposterior direction, were combined by maximum-intensity projection. The resulting images were exported as 16-bit OME-TIFF master files and as 1650 × 960-pixel PNG files for figure preparation.

### Registration to a common anatomical space

Registration was performed using the autofluorescence channel because it provided whole-brain anatomical contrast encompassing the cerebral cortex, hippocampus, striatum, major white-matter tracts and ventral midbrain. Images were first placed in a consistent orientation with dorsal positioned superiorly. For the comparison of molecular markers, the specimen containing *Tac1*, *Pitx2* and *Vglut2* was used as the fixed reference, and the specimen containing TH was registered directly using the corresponding autofluorescence channels. Registration was performed using a Python workflow based on Advanced Normalization Tools through its Python interface, ANTsPyX v0.6.3 (ANTs; RRID:SCR_004757). The procedure comprised global affine alignment followed by nonlinear deformable registration and an additional affine refinement centred on the STN and pSTN region^70,71^. The resulting composite transformation was then applied unchanged to the TH channel.

Transformations were estimated from downsampled 25-50 µm volumes to balance anatomical contrast, computational requirements and registration robustness. The nonlinear deformation field remained anatomically well behaved, with exclusively positive Jacobian determinants (median 1.000; range 0.603–1.651), indicating that the transformation did not introduce folding. Subsequent refinement centred on the STN and pSTN produced modest but consistent improvements in local alignment, increasing intensity normalized cross-correlation from 0.732 to 0.741, edge correlation from 0.320 to 0.344, mutual information from 0.454 to 0.485, and local tissue Dice from 0.9912 to 0.9923. The refinement involved only a small additional correction of approximately 49 µm translation and 0.60° rotation.

Visual assessment of cortical, hippocampal, striatal, white-matter and ventral-midbrain landmarks further supported the regional correspondence. Collectively, these assesments indicated that the registration was suitable for comparing regional signal distributions around the STN and pSTN, while its spatial accuracy was insufficient for interpreting cross-specimen signal overlap as cellular colocalization. Following transformation estimation, the original Level-0 TH data, acquired at a voxel size of 5 × 5 × 10 µm, were resampled directly into the reference brain coordinate system to retain the highest available signal resolution. Custom code was used for multiresolution HDF5/IMS extraction, coordinate conversion, transformation composition and native-resolution resampling. Because TH immunofluorescence and *Tac1* FISH originated from different specimens, overlap between the registered signals was interpreted as regional spatial correspondence within a common anatomical framework and not as evidence of cellular colocalization.

### Serial reconstruction of macaque brain sections

Coronal tissue slabs from the right hemisphere of a macaque monkey brain (10y) were analysed using a custom Python workflow. Twenty-eight (irregularly spaced) DAPI sections were used for structural reconstruction. The marker visualisations contained 24 distinct sections, comprising five *TAC1* FISH and six TH immunofluorescent measurements, with one section containing both markers.

Sections were assembled in a specimen-relative coordinate system. DAPI working images were resampled onto a 50-µm physical-coordinate grid, with background/stripe correction used for structural processing. Initial in-plane poses were obtained from tissue-mask centroids and principal axes. Relative translations between successive available sections separated by no more than eight section numbers were estimated by phase cross-correlation. A bounded, robust least-squares optimisation combined these estimates with pose priors and local serial-smoothness constraints. All sections were visually reviewed for their DAPI orientation. For the final virtual block, the isotropic scale component was removed from the fitted poses to retain scanner-based in-plane scale. These corrected, scale-preserving poses were transferred to marker coordinates, accounting for extraction origins and pixel sampling.

A manually defined rectangular region on, selected via *VGLUT2* overlay to encompass the STN area (7.52 mm by 11.57 mm or 1030 px by 1585 px), was propagated through the existing transforms as fixed block-space regions.

The implemented serial coordinate was calculated from section number using 16 µm per increment (i.e. section thickness), with larger numbers ordered anteriorly.

### Marker specific candidate detection

*TAC1* signal candidates were detected at 3.6496 µm/pixel after per-section percentile contrast scaling. Background-corrected local intensity maxima were characterised using intensity, local contrast, local variance, Laplacian, Hessian-isotropy and gradient features.

TH immunofluorescence was represented as a fluorescence-support field rather than segmented cells. A Gaussian background estimate (σ = 90 pixels) was subtracted, and negative values were set to zero. Analysis was restricted to DAPI-defined tissue at least 35 pixels from its boundary. Reference seeds exceeded the within-section 98.6th percentile of corrected TH intensity. An initial support mask was formed by Gaussian smoothing of the seed mask (σ = 18 pixels), thresholding at 0.008, and five iterations of binary dilation, constrained to the tissue interior. The selected field retained only pixels in this initial mask that also exceeded 0.008 after seed-mask smoothing with σ = 6 pixels and lay within three pixels (approximately 21.9 µm) of a seed. No further dilation was applied. Threshold settings were reviewed with detections or field outlines overlaid on the corresponding native channel images in layered Photoshop documents.

### Three dimensional visualisation and interpretation

Marker coordinates and TH masks were transformed into the common virtual block. The *TAC1*-signal was displayed as red point clouds and the TH-signal as translucent green surfaces generated by marching cubes from the measured section masks.

Final high-resolution TH meshes used the native detection-mask sampling. Thin section surfaces were kept separate across unsampled gaps. Rendering used vedo and VTK with an orthographic camera, fixed framing within each region of interest and a white background. Pooled and individual-section views were exported as 4.096 × 4.096-pixel PNG images. Smooth pooled rotations used 360 independently rendered views at 1° intervals and were encoded using FFmpeg/libx264 as H.264 MP4 files at 30 frames/s. Candidate counts should not be interpreted as independently validated cell numbers and TH fields do not identify individual fibres or TH-positive neurons. Spatial superposition of different serial sections is not interpreted as cellular co-expression.

## Software

The final pipeline used custom Python code with NumPy, SciPy, scikit-image, tifffile, Pillow and psd-tools for processing and review, and vedo/VTK for rendering. Adobe Photoshop was used threshold-overlay review. FFmpeg/libx264 was used for the final MP4 exports.

## Data Availability

Key lab materials used in this study are listed in a Key Resource Table (KRT) alongside their persistent identifiers: https://zenodo.org/records/20007922

All data has been deposited in Zenodo and is found in the same link.

## Code Availability

Custom Python code used for image processing, cross-specimen light-sheet registration, registration quality control, specimen-relative reconstruction of macaque serial sections, quantitative analysis, and visualization is available through GitLab at [https://gitlab.com/asap_mackenzie/dumasnjp2026_3d_rendering_mouse_atlas.git; https://gitlab.com/asap_mackenzie/dumasnjp2026_3d_rendering_atlas.git; https://gitlab.com/asap_mackenzie/DumasNJP2026.git; https://gitlab.com/asap_mackenzie/dumasnjp2026_nhp_rendering.git]. Third-party copyrighted materials from the Paxinos rhesus macaque atlas (including atlas plate images and directly digitized atlas contours or derived meshes) are not included in the repository. Reproduction of the atlas-dependent analyses requires authorized access to the cited atlas and local provision of the required source materials. The repository provides the scripts and instructions needed to process these user-supplied inputs.

## Acknowledgments

The authors thank Dr Anders Björklund, Lund University, Dr Volker Coenen, Freiburg University, San Diego, CA, Dr Nick Hollon, University of California San Diego, CA, Dr Candice Contet, The Scripps Research Institute, San Diego, CA, and Dr Jérôme Baufreton, Université de Bordeaux, for constructive feedback during the course of preparation of this manuscript. The authors also thank Sabrina Leclerc-Turband, Manager at Biobanque Neuro-CEB APHP, France, for post-mortem human brain sections, Nermine Saidi from Institut de la Vision, Paris, France, for slide scanning, and Dr Amanda Schneeweis, Northwestern University, Chicago, IL, for Allen Brain Institute’s MERFISH database guidance.

This research (salaries, experimental costs, mouse housing costs, research facilities) was funded by research funding grants to Å.W.M from the Swedish Research Council (Vetenskapsrådet) [2021-04975], Swedish Brain Foundation (Hjärnfonden) [FO2025-0272-HK-242], the Swedish Parkinson Foundation (Parkinsonfonden) [1632-25][1423-2022], and Hållsten Research Foundation. This research was also funded by Aligning Science Across Parkinson’s [ASAP-020600] [ASAP-027214] through the Michael J. Fox Foundation for Parkinson’s Research (MJFF). For the purpose of open access, the author has applied a CC BY public copyright license to all Author Accepted Manuscripts arising from this submission.

## Author Contributions

Original idea, project administration and funding acquisition, Å.W-M.

Conceptualization, Å.W-M; S.D, A.H.

Methodology, primate brain dissection and tissue block preparation: C.F, C.K.

Methodology, visualization, analysis, and figure preparation (mouse and primate brain section analysis), S.D, A.H, Å.W-M.

Methodology, visualization, analysis, and figure preparation (whole brain analysis, lightsheet microscopy), M.B.R, U.R, A.H.

Methodology, code, brain rendering and analysis: A.H.

Methodology, data-mining (Allen Mouse Brain Atlas dataset (ISH), Allen Brain Cell (ABC) Atlas (MERFISH)) and figure preparation, J.S, Å.W-M.

Contribution of biological material; S.D, C.F, M.B.R, U.R, C.K, Å.W-M.

Writing, original draft, review and editing, Å.W-M.

Writing, review and editing, S.D, A.H.

All authors approved the submitted version of the manuscript.

## Supplementary Video Legends

**Supplementary Video 1. Fluorescent light-sheet microscopy imaging of whole mouse brain in 3D visualizing the detection and spatial distribution of *Pitx2* mRNA.** Raw data coronal fly-through. Abbreviations: pSTN, para-subthalamic nucleus; STN, subthalamic nucleus.

**Supplementary Video 2. Fluorescent light-sheet microscopy imaging of whole mouse brain in 3D visualizing the detection and spatial distribution of *Tac1* mRNA.** Raw data coronal fly-through. Abbreviations: pSTN, para-subthalamic nucleus; STN, subthalamic nucleus.

**Supplementary Video 3. Fluorescent light-sheet microscopy imaging of whole mouse brain in 3D visualizing the detection and spatial distribution of *Pitx2* and *Tac1* mRNAs.** Raw data coronal fly-through. Abbreviations: pSTN, para-subthalamic nucleus; STN, subthalamic nucleus.

**Supplementary Video 4. Specimen-relative three-dimensional point-cloud rendering of *TAC1* mRNA signal in the macaque subthalamic region.** Thresholded *TAC1* fluorescence from serial 16-µm coronal sections is displayed as red points while the reconstruction rotates around the dorsoventral axis.

**Supplementary Video 5. Specimen-relative three-dimensional rendering of the spatial relationship between *TAC1* mRNA signal and TH-immunoreactive fibers in the macaque subthalamic region.** *TAC1* signal is shown in red and TH immunoreactivity in green; their merged distribution illustrates *TAC1* signal within and immediately adjacent to the TH-positive fiber territory along the medial aspect of the STN.

**Supplementary Video 6. Fluorescent light-sheet microscopy imaging of whole mouse brain in 3D visualizing the detection and spatial distribution of *Pitx2* mRNA, mapped to TH immunofluorescence for visualization of dopaminergic cell bodies and projections, including the medial forebrain bundle (MFB).** Coronal fly-through. Abbreviations: pSTN, para-subthalamic nucleus; STN, subthalamic nucleus; *TH*, Tyrosine hydroxylase.

**Supplementary Video 7. Fluorescent light-sheet microscopy imaging of whole mouse brain in 3D visualizing the detection and spatial distribution of *Tac1* mRNA, mapped to *TH* immunofluorescence for visualization of dopaminergic cell bodies and projections, including the medial forebrain bundle (MFB).** Coronal fly-through. Abbreviations: pSTN, para-subthalamic nucleus; STN, subthalamic nucleus; *TH*, Tyrosine hydroxylase.

