## Supplementary figures and images for "Comparative molecular mapping reveals molecular gradients and spatially organized populations in the subthalamic and para-subthalamic regions"

### Suppl Fig 1

Supplementary Fig. 1

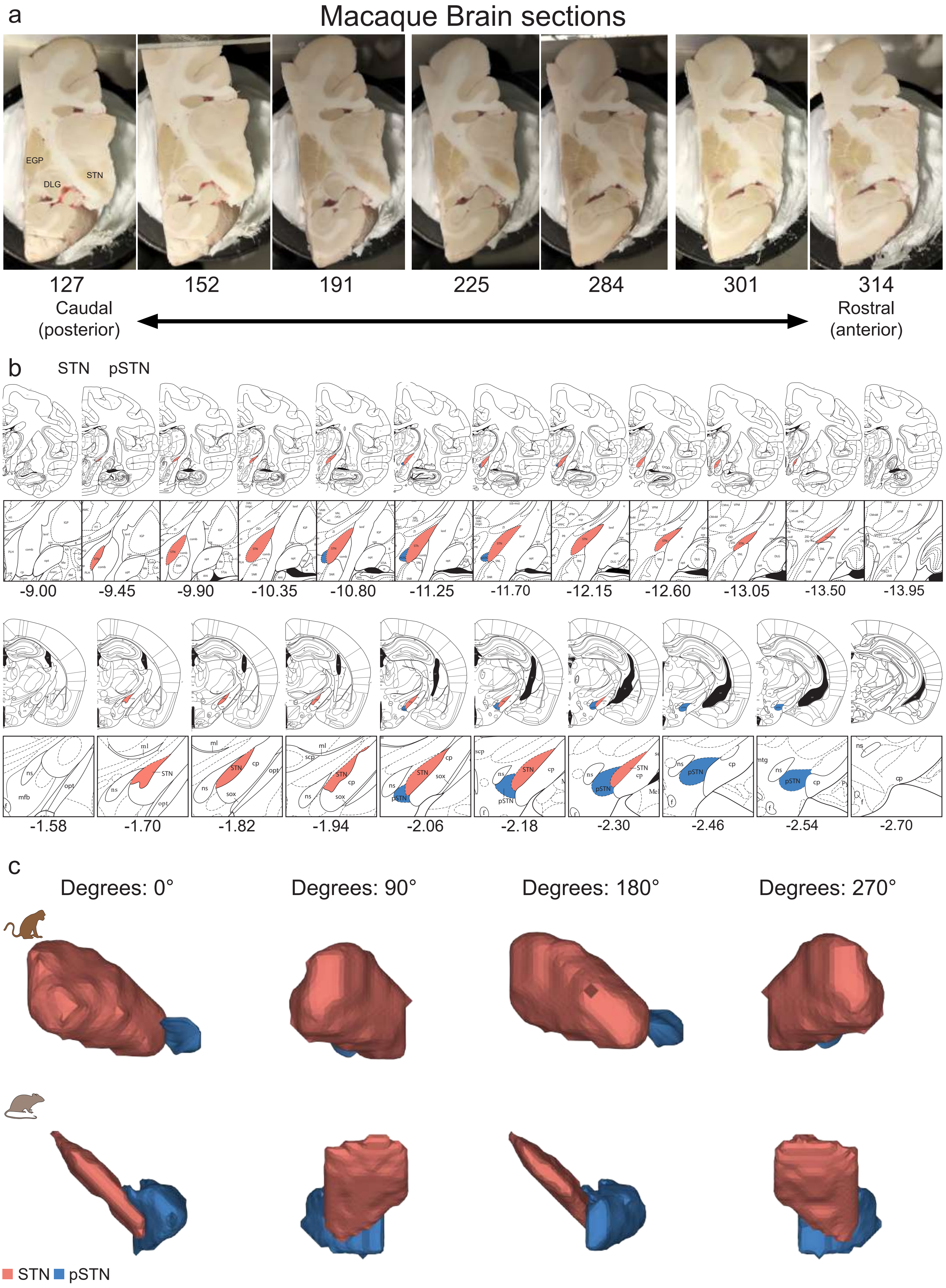

### Suppl Fig 2

**Supplementary Fig. 2**

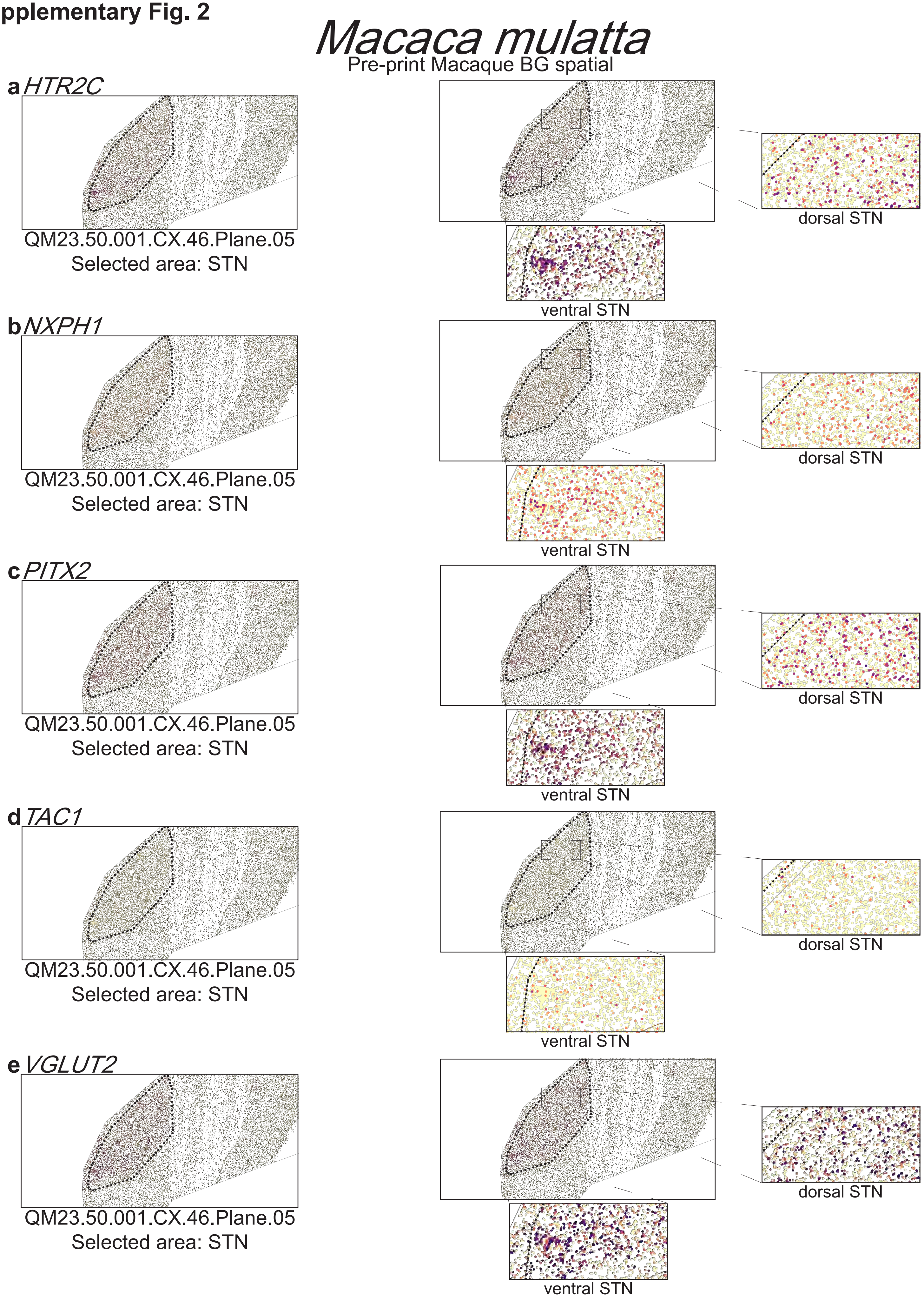

### Suppl Fig 3

**Supplementary Fig. 3**

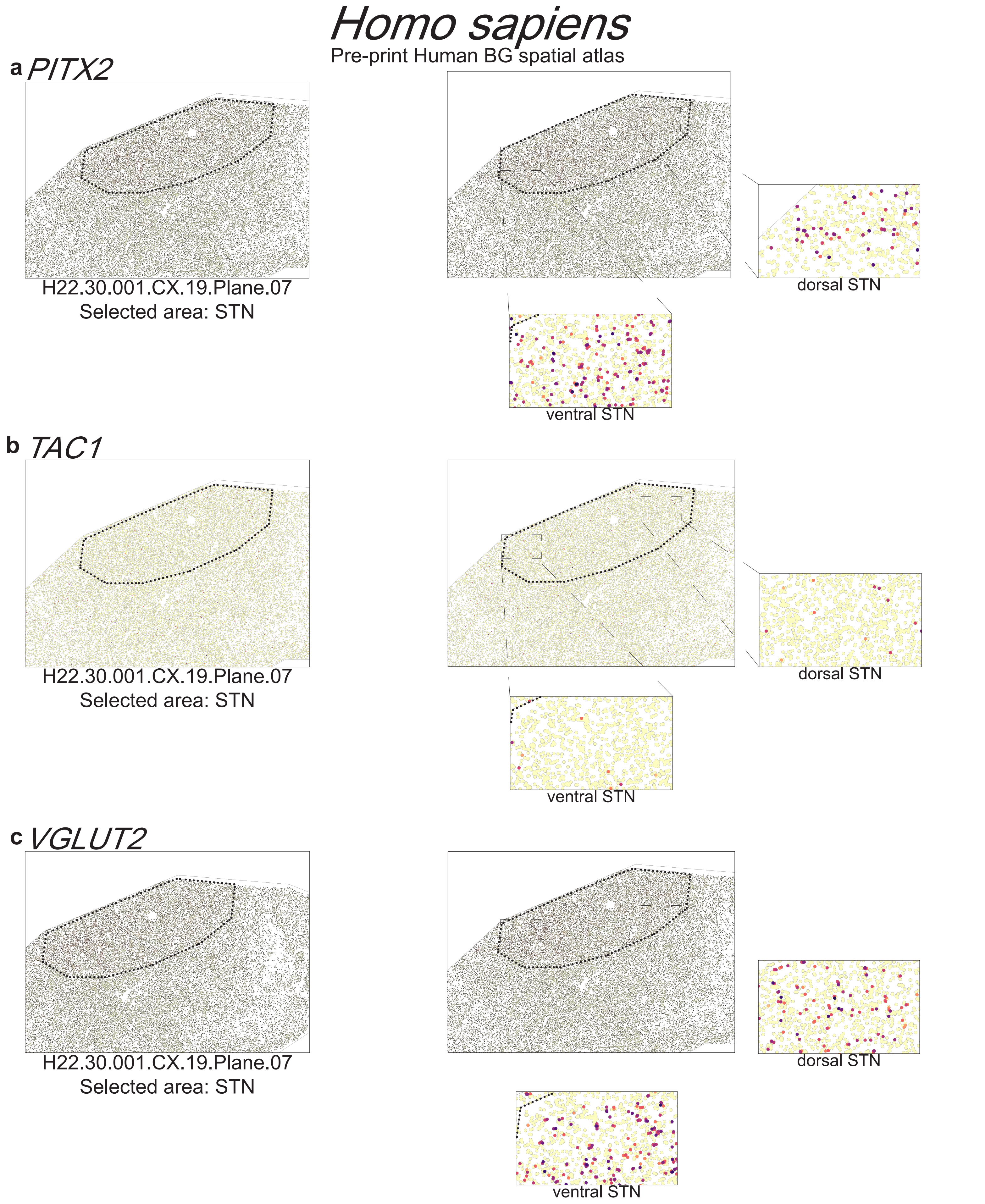

### Suppl Fig 4

Supplementary Fig. 4

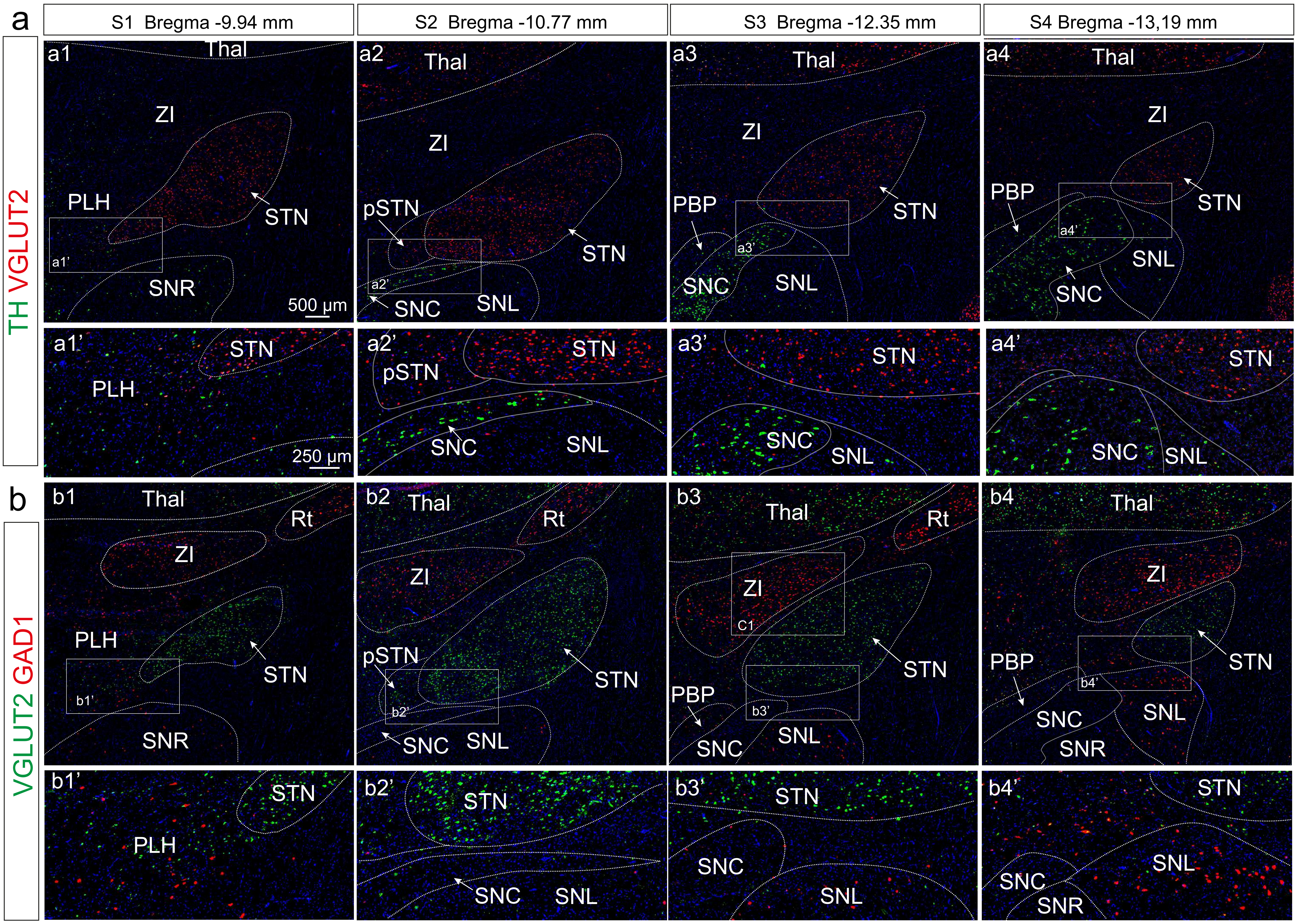

### Suppl Fig 5

#### Supplementary Fig. 5

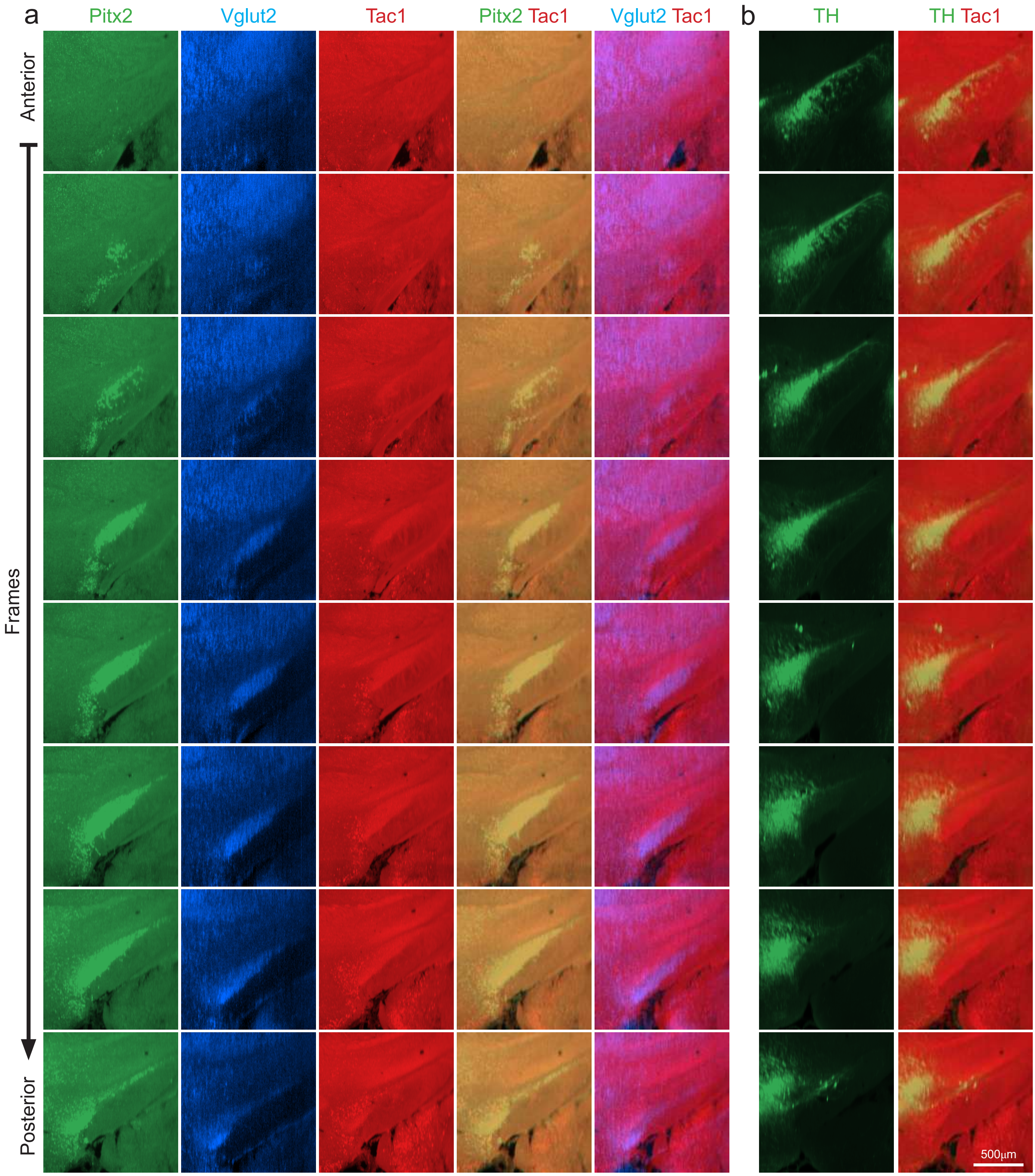

### Suppl Fig 6

Supplementary Fig. 6

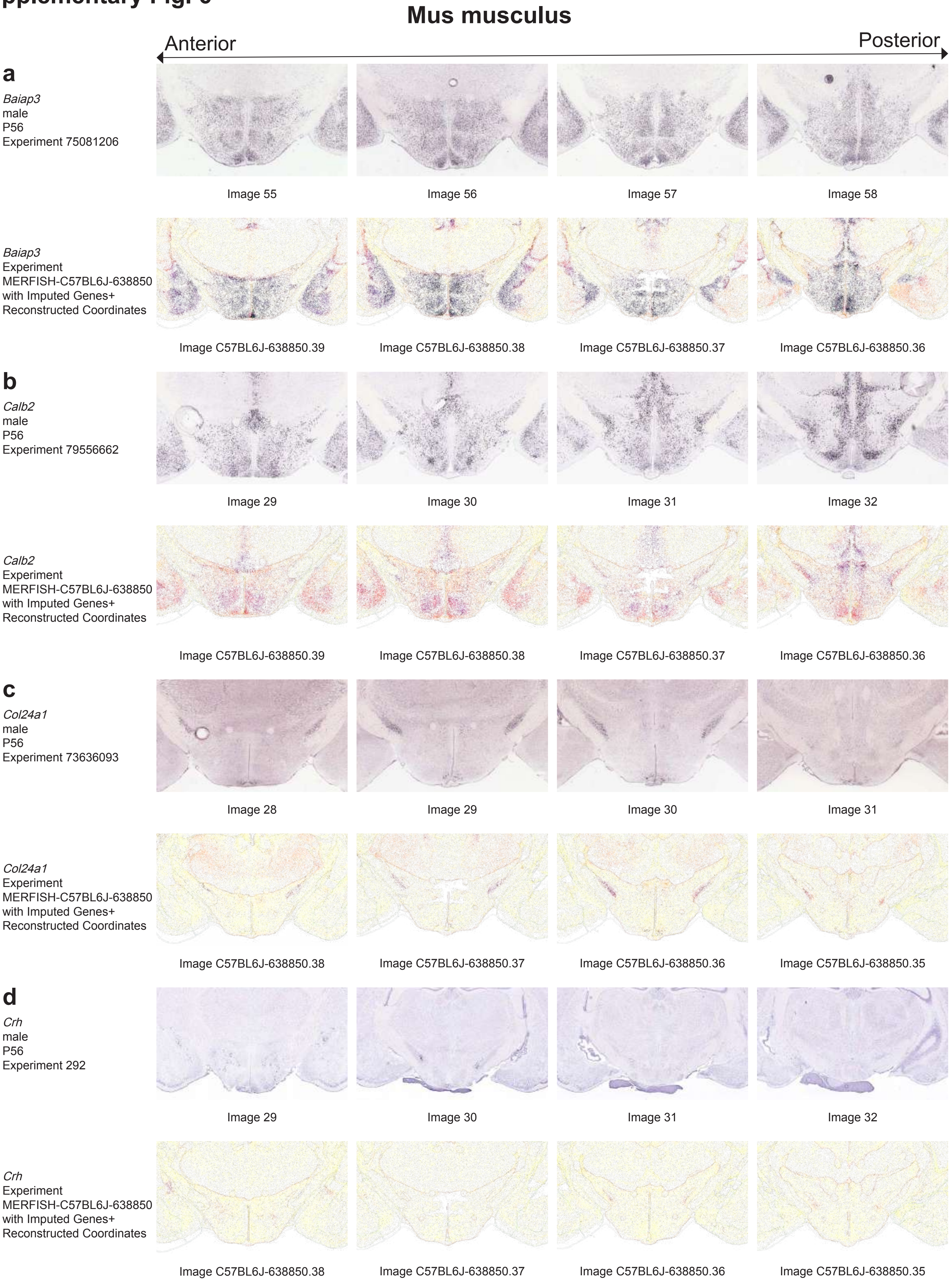

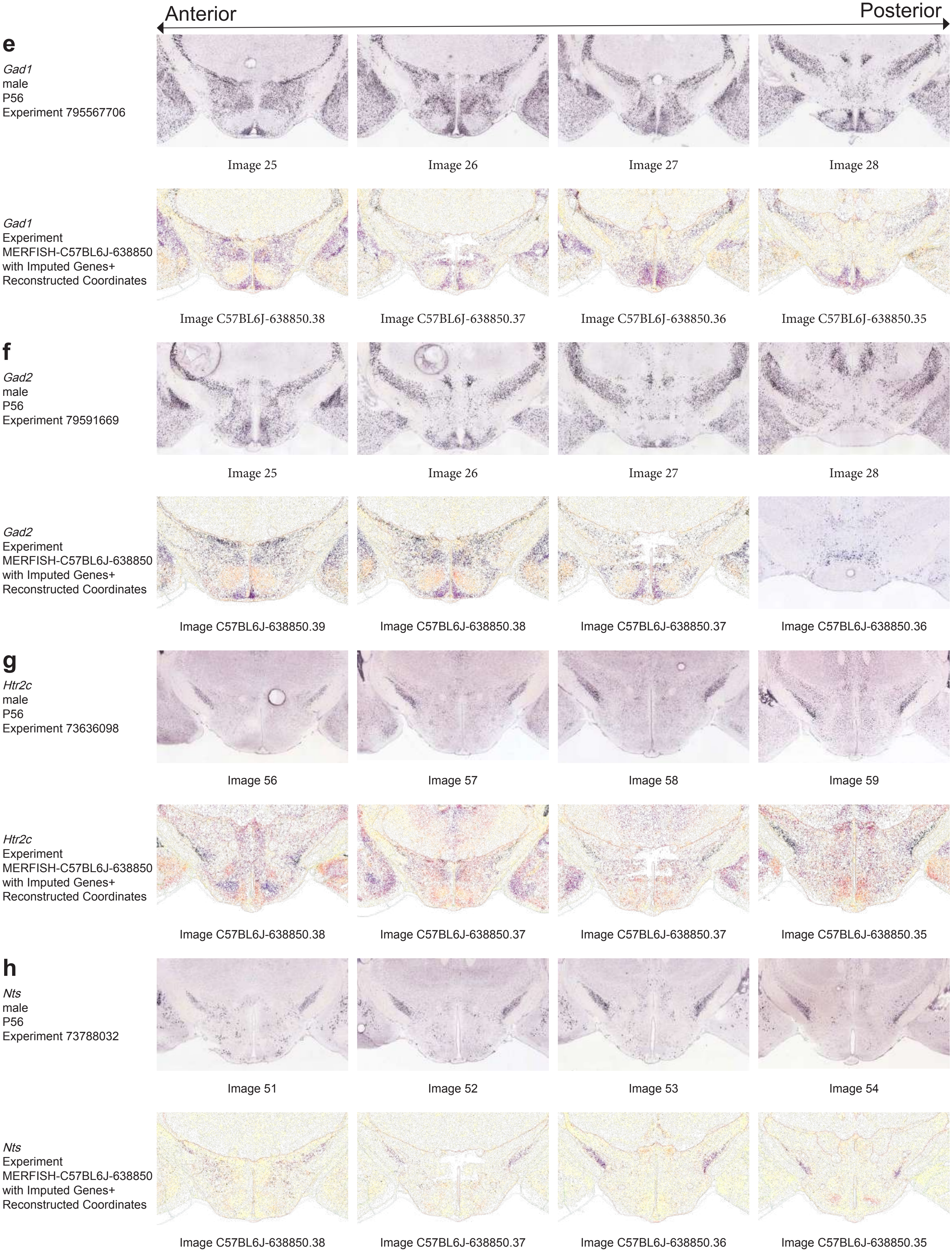

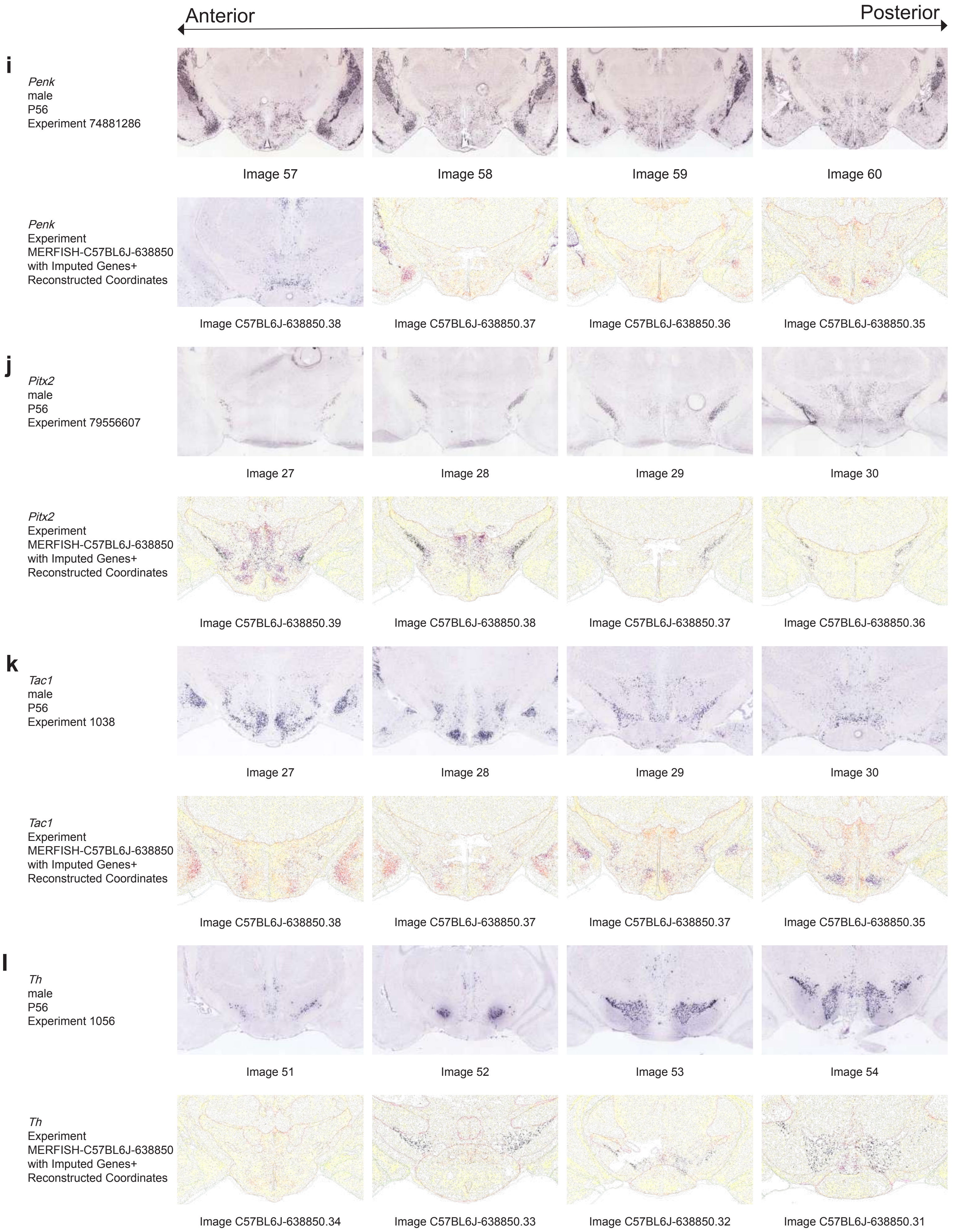

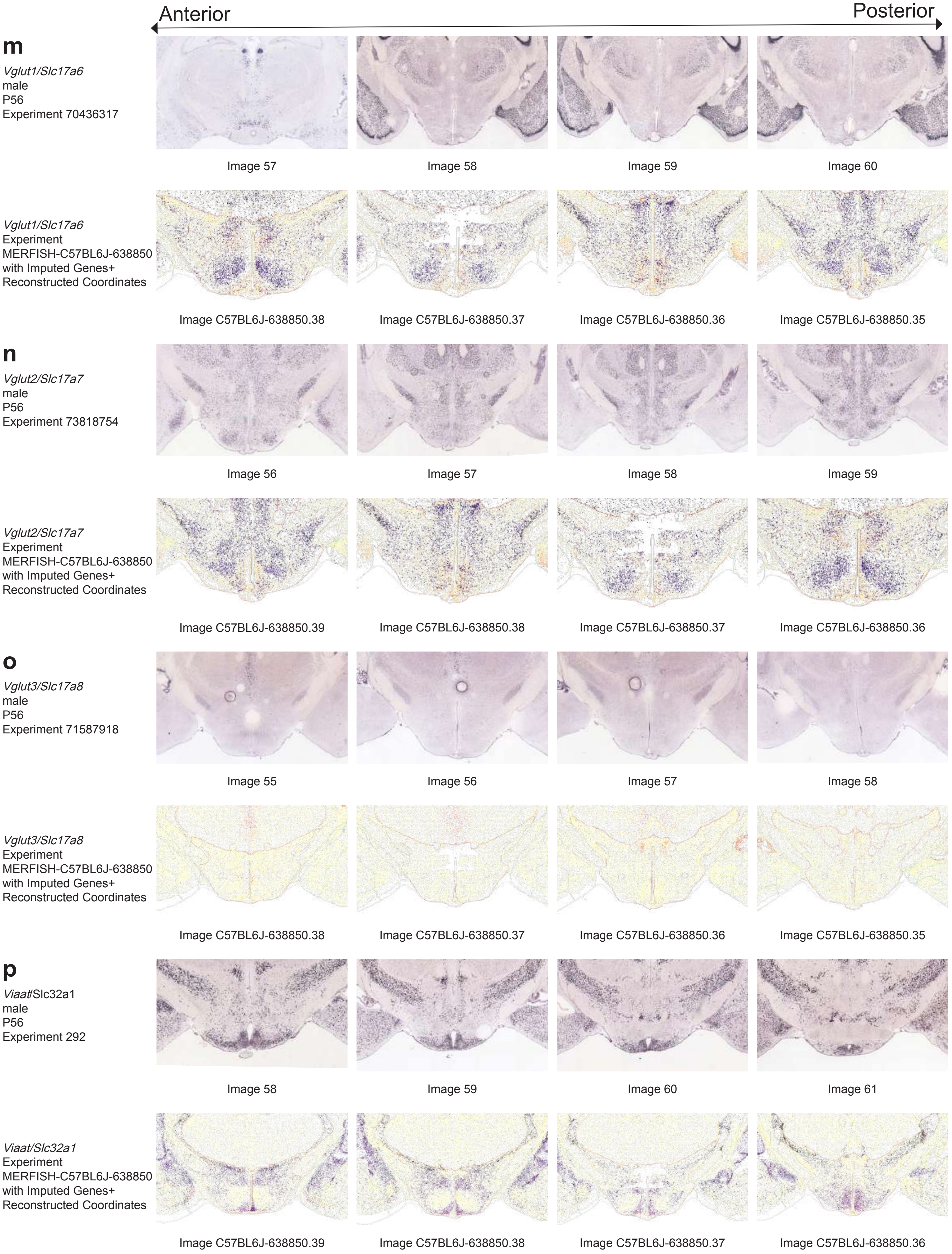

### Suppl Fig 7

Supplementary Fig. 7

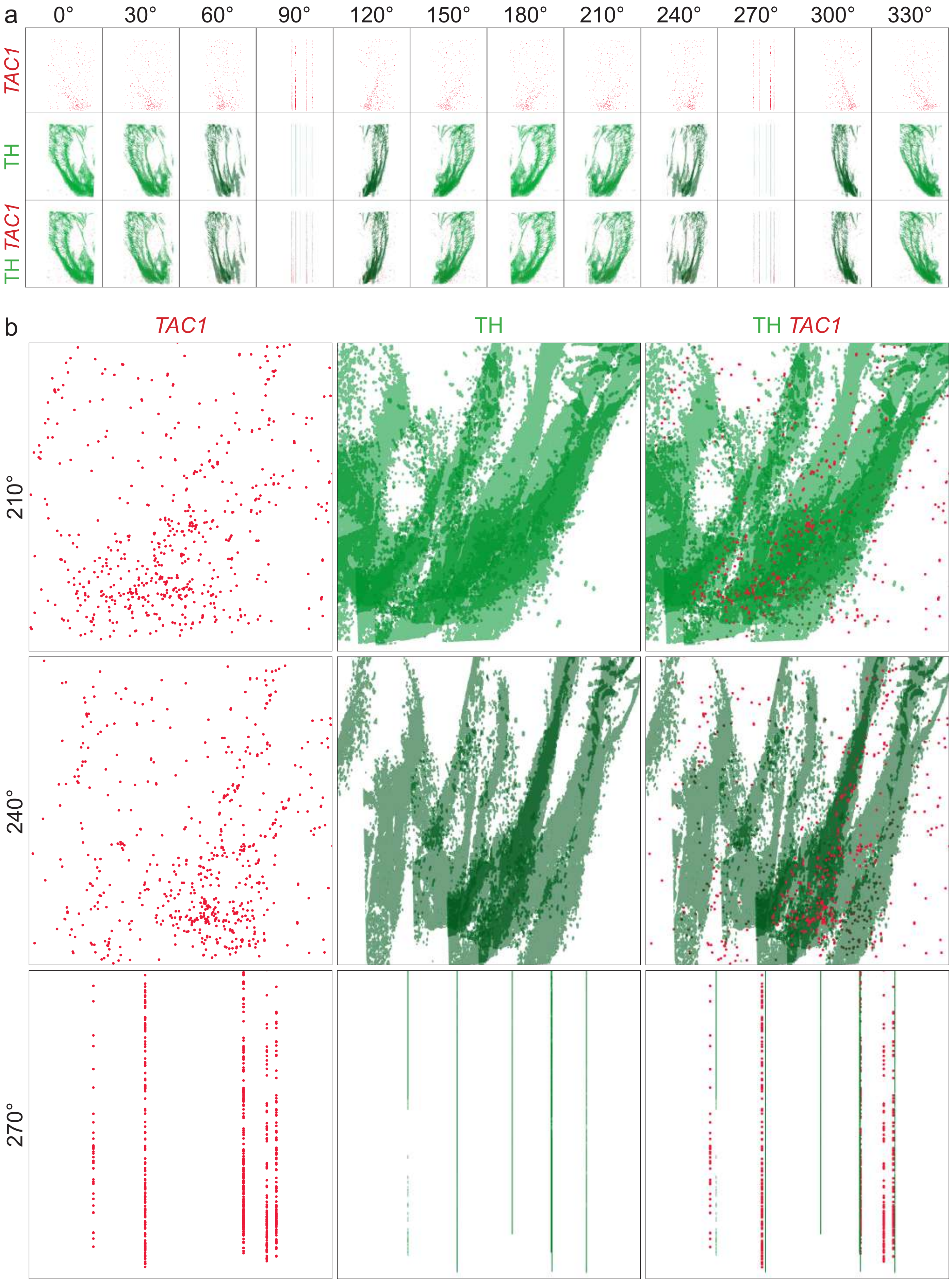

### Suppl Fig 8

#### Supplementary Fig. 8

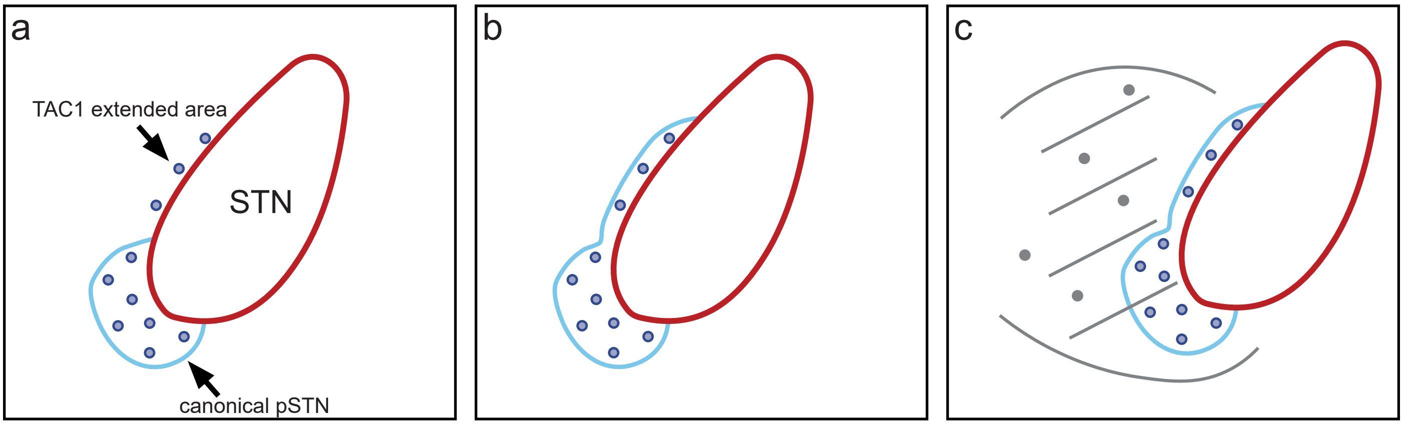
